# MUC13 negatively regulates tight junction proteins and intestinal epithelial barrier integrity via Protein Kinase C

**DOI:** 10.1101/2022.10.27.513982

**Authors:** Celia Segui-Perez, Daphne A.C. Stapels, Ziliang Ma, Jinyi Su, Elsemieke Passchier, Bart Westendorp, Wei Wu, Jos P.M. van Putten, Karin Strijbis

## Abstract

Regulation and adaptation of intestinal epithelial barrier function is essential for human health. The transmembrane mucin MUC13 is an abundant intestinal glycoprotein with important functions for mucosal maintenance that are not yet completely understood. We demonstrate that in intestinal epithelial monolayers MUC13 localized to both the apical surface and the tight junction (TJ) region on the lateral membrane. MUC13 deletion resulted in increased transepithelial resistance (TEER) and reduced translocation of small solutes. TJ proteins including claudins and occludin were highly increased in membrane fractions of MUC13 knockout cells. Removal of the MUC13 cytoplasmic tail (CT) also altered TJ composition but did not result in increased TEER. The increased buildup of TJ complexes in ΔMUC13 and MUC13-ΔCT cells was dependent on PKC, which is in line with a predicted PKC motif in the MUC13 cytoplasmic tail. The responsible PKC member might be PKCδ based on elevated protein levels in the absence of MUC13. Our results identify MUC13 as a central player in TJ complex stability and intestinal barrier permeability.

## Introduction

The intestinal epithelial barrier is a dynamic system that prevents bacterial invasion while at the same time allowing the transport of nutrients (*1, 2*). The intestinal mucosal epithelium consists of various types of enterocytes and a closely associated mucus layer in which highly glycosylated mucin proteins are the main structural component. Mucins can be categorized into soluble mucins that are secreted by goblet cells and transmembrane (TM) mucins that are cell-bound and expressed by most types of enterocytes. TM mucins expressed in the human intestinal tract include MUC1, MUC3, MUC12, MUC13, and MUC17 (*3*) of which MUC13 shows the most widespread expression along the different segments of the gastrointestinal tract (*4*). The extracellular domains of TM mucins are highly glycosylated and their cytoplasmic tails have signaling capacity (*2*). TM mucins are highly diverse, and the different members have been implicated in fundamental epithelial processes including the regulation of cell-cell interactions, proliferation, differentiation, apoptosis, and modulation of inflammatory responses (*2, 5, 6*). Dysfunction of TM mucins has been associated with the development of inflammatory bowel disease (IBD) including ulcerative colitis (UC) and Crohn’s disease (CD) (*7*–*9*). Reduced intestinal barrier function and the translocation of bacterial components across the intestinal mucosal-epithelial barrier are hallmarks of IBD. The contributions of specific TM mucins to epithelial barrier integrity and development of IBD remain to be established.

MUC13 is a relatively small TM mucin that consists of a glycosylated extracellular domain (ED) that contains a Sperm protein, Enterokinase, and Agrin (SEA) domain, three epithelial growth factor (EGF)-like domains, and a cytoplasmic tail (CT) with putative phosphorylation sites. Previous studies demonstrated MUC13 expression on the apical surface of polarized epithelial cells, and cytoplasmic and nuclear localization was observed in colorectal cancer (CRC) and during metastasis (*4, 10*). MUC13 mRNA expression is upregulated in the inflamed colon in IBD patients (*11*) and a mutation in the MUC13 cytoplasmic tail was shown to be associated with the development of UC (*12, 13*).

The function of MUC13 seems to be multifaceted as it has been linked to different aspects of mucosal maintenance and inflammation. Overall, most MUC13-associated phenotypes can be considered pro-inflammatory and promote wound healing and tumorigenesis. MUC13 enhances the epithelial pro-inflammatory response to bacterial ligands (*14*) and interacts with tumor necrosis factor receptor 1 (TNFR1) thereby promoting TNF-induced NF-kB activation (*15*). Muc13-deficient mice and human intestinal MUC13 knockdown cells are more sensitive to toxin-induced apoptosis (*11*). Single-cell migration is enhanced in colon cancer cells with MUC13 overexpression (*16*). In pancreatic ductal adenocarcinoma (PDAC) cells, MUC13 interacts with HER2 resulting in activation and cytoskeletal remodeling, growth, motility, and invasive growth (*17*). Thus, MUC13 seems to be a key protein that is linked to several aspects of intestinal epithelial health and disease, but the underlying molecular mechanisms remain to be resolved.

Epithelial barrier integrity is critically regulated by the junction complexes that are embedded in the lateral membranes of neighboring cells. The junction complexes can be divided into adherence junctions (AJ), tight junctions (TJ), and desmosomes. Together, they form the apical junctional complex which seals the paracellular space between cells (*18*). TJ are large multimeric protein complexes in the lateral membrane that consist of various transmembrane proteins, including occludin and claudins (*19, 20*). The main function of TJs is the regulation of paracellular permeability, but they also play a role in polarization, morphogenesis, cell proliferation, and regulation of gene expression (*21*). Intracellularly, proteins such as ZO-1 connect the TJ complex to the actin cytoskeleton and signal transduction molecules (*22, 23*). AJ and desmosomes are present along the full length of the lateral membrane, connecting adjacent cells, and contribute to the barrier function without sealing the paracellular space (*19*). The main structural protein of AJs is E-cadherin. Through its intracellular tail, E-cadherin interacts with β-catenin, the central regulator of the epithelial WNT pathway (*24*). Changes in barrier function and TJ and AJ proteins are often observed in IBD (*25*–*28*).

Multiple members of the TM mucin family have been implicated in the regulation of cell-cell interactions. MUC1, MUC4, and MUC16 all reduce the interaction between E-cadherin and β-catenin at the membrane, thereby promoting β-catenin translocation to the nucleus and subsequent activation of the Wnt signaling pathway (*29*–*33*). MUC1 and MUC16 can interact directly with β-catenin via the phosphorylated cytoplasmic tail (*34, 35*), whereas MUC13 can enhance nuclear translocation of β-catenin through interaction with GSK-3β (*36*). Several studies have linked MUC1, MUC16, and MUC17 with alterations in TJ proteins, thereby influencing epithelial monolayer properties, though the underlying mechanisms are not yet understood (*37*–*40*). Whether MUC13 regulates TJ proteins and epithelial barrier integrity is yet unknown.

In the present study, we investigate the function of MUC13 in the regulation of barrier integrity of the intestinal epithelium. Our data identify MUC13 as a central regulator of tight junction strength and paracellular passage which has important implications for the role of this TM mucin in IBD and colorectal cancer development.

## Results

### MUC13 is highly expressed in the intestinal tract and localizes to the apical and lateral membrane

To determine the expression of MUC13 and other mucin genes in different segments and cell types of the gastrointestinal tract, we analyzed the gut atlas single-cell RNA-sequencing dataset (gutcellatlas.org). This dataset contains 428,000 intestinal cells from fetal, pediatric, and adult donors. We focused on the adult cells and extracted the average expression of the different transmembrane and soluble mucins from each part of the gastrointestinal tract. MUC13 was expressed in at least 50% of the cells across all locations (Fig. 1A). MUC3A was also detected in all segments but the expression was lower in the appendix and rectum. MUC1 and MUC4 expression was mainly observed in the colon and rectum, while MUC17 showed the opposite pattern with high expression in the small intestine. We then analyzed the dataset for mucin expression within different cell type lineages. As expected, high expression of the secreted mucin MUC2 was observed for goblet cells. Transmembrane mucins MUC1 and MUC4 were also highly expressed in goblet cells. A comparable expression pattern was found for MUC13 and MUC3A with high expression throughout all cell types with the highest levels in enterocytes and BEST4+ epithelial cells (Fig. 1B).

**Fig. 1.**
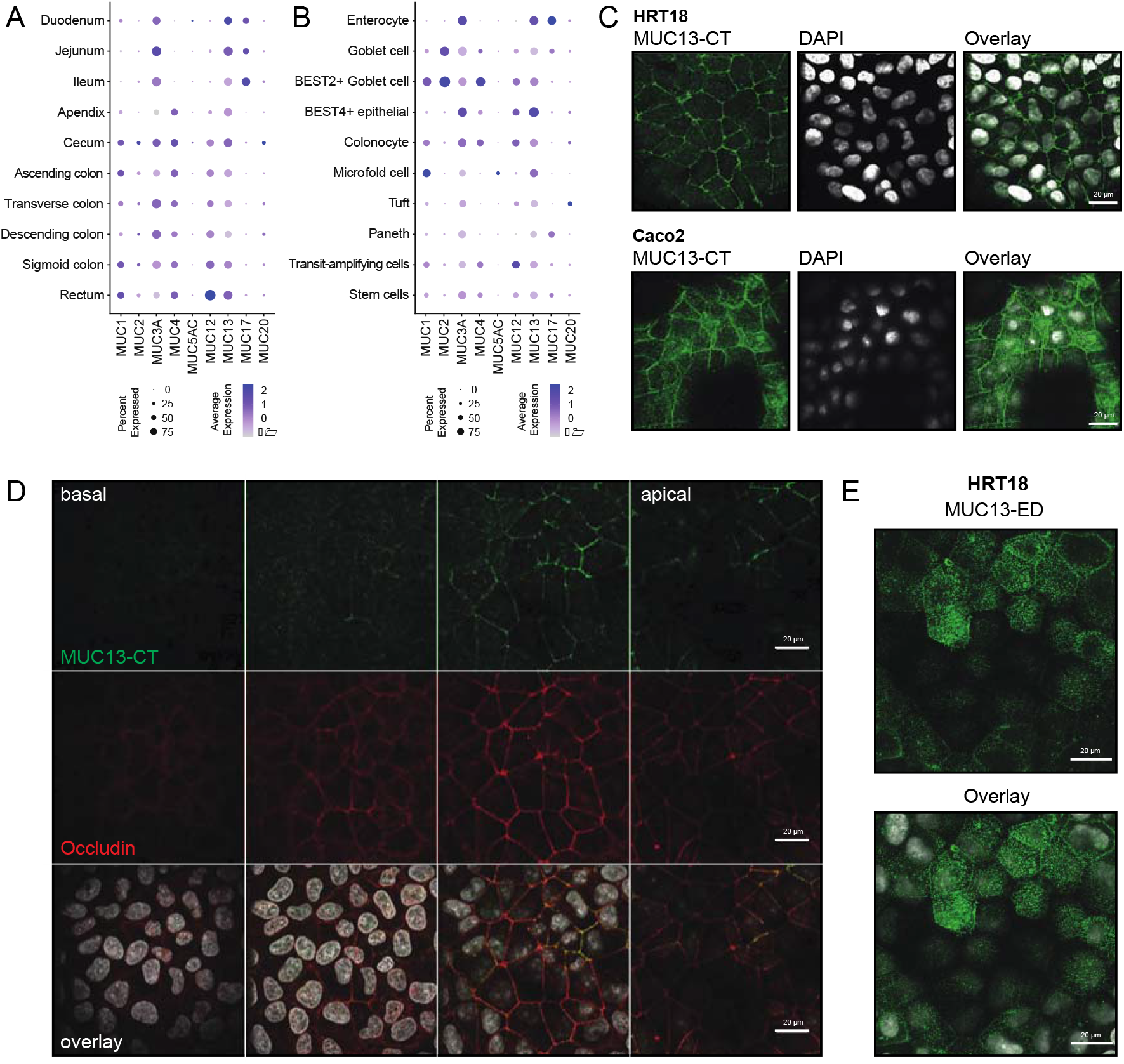
MUC13 is highly expressed in the intestinal epithelium and localizes at the apical and lateral membrane. (**A-B**) Single-cell RNA-sequencing data of adult donors showing expression levels of mucin genes along each section of the intestinal tract (**A**) and by different cell types (**B**). (**C**) Immunofluorescence microscopy of HRT18 and Caco-2 intestinal cells stained for MUC13 cytoplasmic tail (MUC13-CT) (green) and nuclei (white). (**D**) Immunofluorescence microscopy of HRT18 cells with antibodies against MUC13-CT and occludin, in combination with DAPI from basal to lateral Z planes. (**E**) Immunofluorescence microscopy of HRT18 cells with monoclonal MUC13 antibody against the extracellular domain. White scale bars represent 20 mM.

MUC13 has been reported to localize to the apical surface of differentiated intestinal epithelial tissue (*4, 10*). We determined the expression and localization of MUC13 in intestinal epithelial HRT18 and Caco-2 cells. Immunofluorescence confocal microscopy was performed with a MUC13 antibody directed against the cytoplasmic tail. With this antibody, the majority of MUC13 was detected on the lateral membranes of both HRT18 and Caco-2 cells (Fig. 1C). By creating Z-stacks, we observed MUC13 staining from the apical side of the lateral membrane towards the middle and limited or no staining in the basal planes which depict the lower region of the lateral membrane. The tight junction protein occludin also localized to the top half of the lateral membrane, similar to MUC13 (Fig. 1D). Staining of the adherence junction protein E-cadherin was observed along the entire lateral membrane (Fig. S1). Using a previously described method for transmembrane proteins (*41*), we generated a novel monoclonal antibody against the extracellular domain of MUC13. This antibody stained both the apical surface and upper part of the lateral membrane in HRT18 cells (Fig. 1E). These results demonstrate that different MUC13 antibodies recognize distinct MUC13 populations on the apical and lateral membranes. We conclude that MUC13 localizes to the apical surface of enterocytes and the apical side of the lateral membrane in the region where tight junctions are found.

### Deletion of MUC13 and targeted deletion of the MUC13 cytoplasmic tail using CRISPR/Cas9

To study the function of the full-length MUC13 protein and the contribution of the MUC13 cytoplasmic tail, we designed CRISPR/Cas9 strategies to generate two types of HRT18 MUC13 knockout cell lines. Expression of the full-length MUC13 protein was eliminated by deletion of 380 base pairs in the second exon which resulted in disruption of the reading frame (Fig. 2A). As a control, HRT18 cells were transduced with an empty CRISPR plasmid without guide RNAs, and the resulting cell line was used in all the experiments as accompanying wild type (HRT18-WT). For targeted removal of the MUC13 cytoplasmic tail, we selected gRNAs that target exon 10 and were predicted to result in the removal of 121 bp that encode the majority of the MUC13 cytoplasmic tail (Fig. 2A). For all genotypes, we generated two independent cell lines resulting in two HRT18-WT (WT 1 and 2), two HRT18-ΔMUC13 (ΔMUC13 1 and 2), and two HRT18-MUC13-ΔCT cell lines (MUC13-ΔCT 1 and 2). The domain structures of the MUC13 WT and ΔCT proteins are depicted in Fig. 2B, and the amino acid sequence of each domain is shown in Fig. 2C. The resulting deletion and disrupted reading frame of the different cell lines were confirmed by PCR and sequencing (Fig. 2D). The two ΔMUC13 cell lines lacked 300 and 377 bp fragments, respectively. Both MUC13-ΔCT clones had a deletion of 121 bp resulting in a stop codon three amino acids after the deletion. The predicted sequence of the remaining cytoplasmic tail is ARSNNKTKHIEEENLIDEDFQNLKLRSIR*, which lacks multiple putative phosphorylation sites and a predicted PKC motif that are present in the full-length cytoplasmic tail.

**Fig. 2.**
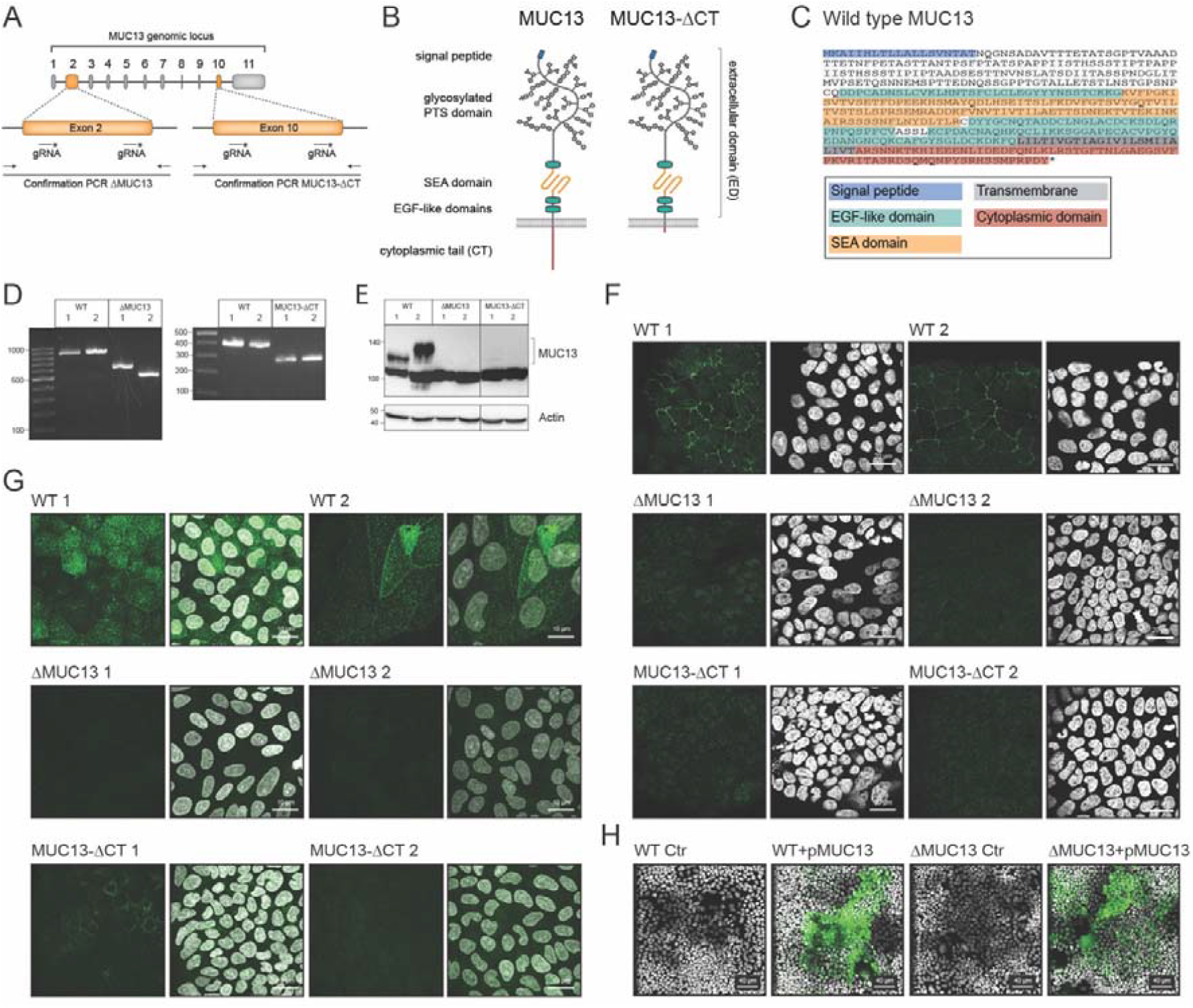
Generation of MUC13 knockout and MUC13-GFP overexpression cell lines. (**A**) CRISPR/Cas9 targeting strategy using two guide RNAs directed against exon 2 or exon 10 of MUC13. (**B**) Schematic representation of WT and MUC13-ΔCT MUC13 domain structure. (**C**) Wild type MUC13 protein sequence with domains color-coded as in Fig. 2B. (**D**) Confirmation PCR of WT and ΔMUC13 cell lines (left), and WT and MUC13-ΔCT cell lines (right). (**E**) Immunoblot of WT, ΔMUC13, and MUC13-ΔCT cell lines with anti-MUC13-CT antibody and actin loading control. Molecular mass standards (kDa) are indicated on the left. (**F)** Immunofluorescence confocal image of WT, ΔMUC13, and MUC13-ΔCT cells stained for MUC13-CT (green) and nuclei (white). White scale bars represent 20 mM. (**G**) Immunofluorescence confocal images of WT, ΔMUC13, and MUC13-ΔCT cells stained for MUC13-ED (green) and nuclei (white). White scale bars represent 10 mM. (**H**) Immunofluorescence confocal image of WT Ctr (empty plasmid), WT+pMUC13 (with inducible MUC13-GFP construct), ΔMUC13 Ctr, and ΔMUC13+pMUC13 complementation cell lines after doxycycline induction for 24h. MUC13-GFP is depicted in green, and nuclei are shown in white. White scale bars represent 40 mM.

Next, we investigated the expression of MUC13 or the truncated protein in HRT18-WT, ΔMUC13, and MUC13-ΔCT cells. Western blot analysis with the MUC13-CT antibody showed MUC13-reactive bands of 120 kDa and 130 kDa in the WT cell lines, which were absent in ΔMUC13 and MUC13-ΔCT cells (Fig. 2E). A slightly different molecular weight was observed for MUC13 in the two WT cell lines which could be the result of differential glycosylation or processing and/or activation by (auto)proteolytic cleavage as has been reported for MUC1 (*2, 42, 43*). MUC13 expression in the different cell lines was also investigated by confocal microscopy. With the antibody directed against the cytoplasmic tail, we observed lateral membrane staining in HRT18-WT cells, while the signal was absent in ΔMUC13 and MUC13-ΔCT cells (Fig. 2F). With the antibody directed against the extracellular domain, we observed apical and lateral staining in WT cells, but not in ΔMUC13 cells. For the MUC13-ΔCT 1 cell line, we noted reduced intensity staining of the extracellular domain compared to the WT cells, while the extracellular domain was barely detectible in the MUC13-ΔCT 2 cell line (Fig. 2G). These results demonstrate that while they are genetically identical, expression levels differ between the two ΔCT cell lines which might be due to reduced stability of the truncated MUC13 protein. We conclude that our CRISPR-Cas9 strategy in the intestinal epithelial HRT18 cells was successful and resulted in MUC13 knockout cell lines and cell lines that express MUC13 without the cytoplasmic tail.

### Generation of MUC13-GFP overexpression and complementation cell lines

To complement MUC13 in the knockout cell lines, we cloned a doxycycline-inducible MUC13-GFP plasmid with a codon-optimized MUC13 DNA sequence that left the amino acid sequence unaltered but allowed cloning and expression. Lentiviral transduction was used to introduce the MUC13-GFP construct into HRT18 wild type and ΔMUC13 cells resulting in overexpression WT+pMUC13 and complemented ΔMUC13+pMUC13 cell lines. Doxycycline induction resulted in MUC13-GFP expression in at least 50% of the total cell populations (Fig. 2H).

### Deletion of MUC13 alters epithelial barrier properties

To investigate the contribution of MUC13 to epithelial barrier properties, we grew the HRT18 cell lines for two weeks to allow the buildup of cell junctions. Cells reached full confluency on day 3. To determine the monolayer architecture, we performed immunofluorescence microscopy and stained for occludin, E-cadherin, and nuclei (DAPI). All cell lines formed confluent monolayers with comparable occludin and E-cadherin staining. ΔMUC13 cells did show a less rounded cell morphology compared to WT and MUC13-ΔCT cells (Fig. 3A). Next, confluent monolayers were grown on membranes in Transwell plates and differentiated for 14 days. Transepithelial electrical resistance (TEER), a measure of TJ strength based on electrical resistance, was determined over time for all cell lines. The TEER of ΔMUC13 clones was, on average, three-times higher compared to WT cells while the TEER of the MUC13-ΔCT cells was comparable to WT cells (Fig. 3B, C). To rule out the possibility that differences in TEER were caused by differences in cell numbers, we counted the number of nuclei per plane after 14 days of differentiation. The numbers of nuclei were comparable between the cell lines, indicating that the difference in TEER was not a result of differences in proliferation (Fig. 3D). Next, we determined the buildup of TEER in the MUC13 overexpression and complementation cell lines WT+pMUC13 and ΔMUC13+pMUC13. Overexpression of MUC13-GFP in the ΔMUC13 background led to a significant reduction of TEER buildup over time, while overexpression in the wild type background did not affect TEER (Fig. 3E, F). Together, these data indicate that MUC13 negatively regulates TEER buildup.

**Fig. 3.**
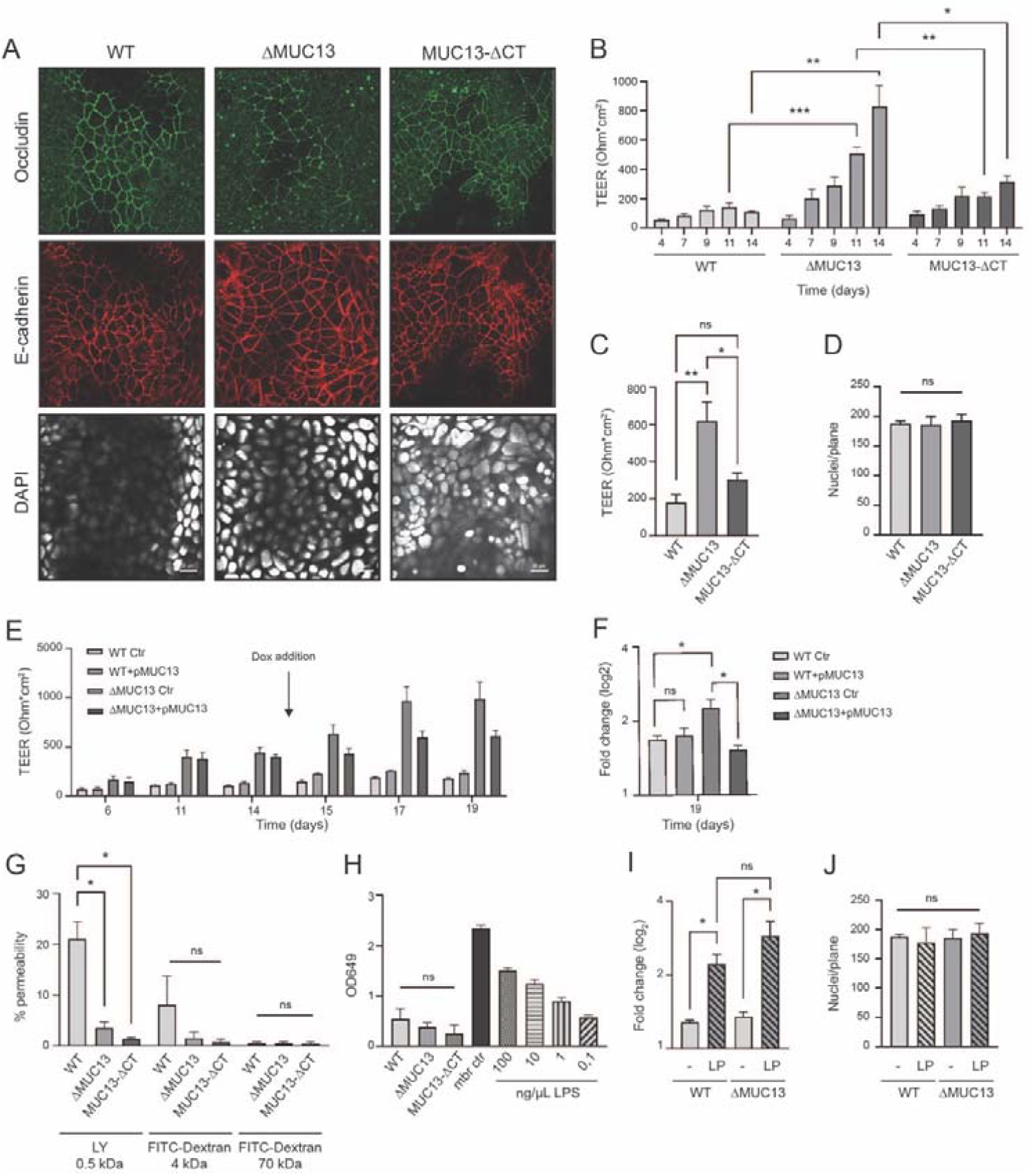
Deletion of MUC13 alters epithelial barrier properties. (**A**) Immunofluorescence confocal image of WT, ΔMUC13, and MUC13-ΔCT cell monolayers showing occludin (green), E-cadherin (red), and nuclei (DAPI; white) staining. White scale bars represent 20 mM. (**B**) Transepithelial electrical resistance (TEER) buildup in cell monolayers grown for up to 14 days. (**C**) TEER buildup in 2-weeks-differentiated monolayers. (**D**) Quantification of cell nuclei per plane by confocal microscopy (DAPI) in cell monolayers after 14 days of differentiation. (**E**) TEER buildup in the MUC13 overexpression and complementation WT+pMUC13 and ΔMUC13+pMUC13 cell lines. Doxycycline was added on day 14 as indicated by an arrow. (**F**) Fold change (log2) of TEER increase in WT+pMUC13 and ΔMUC13+pMUC13 cells on day 19 compared to day 14 before the addition of doxycycline. (**G**) Paracellular passage of Lucifer Yellow CH substrate and FITC-dextran particles in 14-days-differentiated cell monolayers. (**H**) Paracellular permeability assay with LPS from *Escherichia coli* 0111:B4 in 14-days differentiated monolayers. (**I**) Fold change (log2) compared to time 0 of TEER increase in 14 days-differentiated WT and ΔMUC13 cell monolayers after addition of *Lactobacillus plantarum* (LP) for 42 h at MOI 50. (**J**) Quantification of cell nuclei per plane by confocal microscopy (DAPI) in WT and ΔMUC13 cell monolayers after 42 h incubation with LP. All graphs represent the average and SEM of three independent experiments. ns, non-significant; *, p<0.05; ** p<0.01; *** p<0.001.

### MUC13 deletion leads to decreased paracellular passage of small molecules

TEER reflects the conductance of small ions via the paracellular pathway, which represents the passage of molecules through the intercellular spaces between adjacent epithelial cells. The flux of larger molecules through the paracellular pathways can be addressed using organic tracers, such as Lucifer Yellow CH and fluoresceinated (FITC)-dextran particles. We seeded our cell lines on Transwell membranes as before and the transfer of compounds from the apical compartment to the basolateral side was determined. WT, ΔMUC13, and MUC13-ΔCT cells were all highly restrictive for the passage of 4 and 70 kDa FITC-dextran particles. For the smaller 520 Da Lucifer Yellow tracer, ΔMUC13, and MUC13-ΔCT monolayers were restrictive while WT cells were permeable (Fig. 3G). Because translocation of bacterial endotoxin lipopolysaccharide (LPS) across the intestinal barrier is an important hallmark of intestinal barrier dysfunction, we determined the passage of *Escherichia coli* 0111:B4 lipopolysaccharide (LPS-EB). A maximum of ∼300 ng/μL LPS reached the basal compartment after 24 hours incubation for the control wells, and less than 0.1 ng/μL LPS passage for the different HRT18 cell lines. LPS passage was comparable between WT, ΔMUC13, and MUC13-ΔCT cell lines indicating the restrictiveness of these cells to the passage of larger particles (Fig. 3H). In summary, we observed that deletion of MUC13 results in a higher buildup of TEER and lower paracellular passage of the small organic solute Lucifer Yellow compared to WT. The TEER of the MUC13-ΔCT cell line was comparable to WT, but a significant restriction of Lucifer Yellow passage compared to WT was observed. We conclude that the paracellular pathway is altered in both MUC13 deletion cell lines.

### Epithelial barrier strengthening by *Lactobacillus plantarum* is independent of MUC13

To investigate the role of MUC13 in TEER regulation, we made use of a probiotic bacterium known to enhance TEER formation. *Lactobacillus plantarum* (LP) is a commensal of the large intestine that can enhance intestinal barrier function by activating Toll-like receptor 2 (TLR2) signaling which triggers the translocation of occludin and ZO-1 to the TJ region in Caco-2 cells (*44, 45*). LP was added to 14 days-differentiated WT and ΔMUC13 cell monolayers at a multiplicity of infection (MOI) of 50. TEER was measured every 12 h for two days and the largest increase was observed 42 h after the addition of the bacteria. The incubation with LP resulted in a comparable increase in TEER in WT and ΔMUC13 cells by 2.2-fold and 2.8-fold, respectively (Fig. 3I). To confirm that the increase in TEER was not due to increased cell counts we confirmed that the number of nuclei per plane at 42 h post-infection was comparable between the different cell lines and conditions (Fig. 3J). These data show that the increase in epithelial barrier properties in response to LP is not dependent on MUC13. The function of MUC13 in epithelial barrier regulation seems therefore not linked to the pathway of TLR-mediated increase of ZOs and occludin proteins induced by LP.

### Tight junction proteins are highly upregulated in the absence of MUC13

Based on the TEER and translocation data, we hypothesized that ΔMUC13 cells might have stronger tight junctions that reduce the paracellular translocation of ions and small particles. The functional fraction of junction proteins localizes to the plasma membrane. To be able to detect this functional fraction with increased sensitivity, we developed a fractionation protocol to enrich for the membrane fractions of WT, ΔMUC13, and MUC13-ΔCT monolayers (Fig. 4A). We verified the fractionation strategy by Western blot with Na^+^/K^+^-ATPase (as membrane marker), Histone-H3 (as nuclear marker), and β-actin (as cytoplasmic marker). Membrane fractions showed a successful enrichment of membrane proteins and were free of nuclear contamination (Fig. 4B). Next to the validation by Western blotting, we further analyzed the membrane fractions by mass spectrometry. From three technical replicates, a total of 4.054 proteins were identified, of which 3.916 proteins were quantifiable in at least two out of three replicates.

**Fig. 4.**
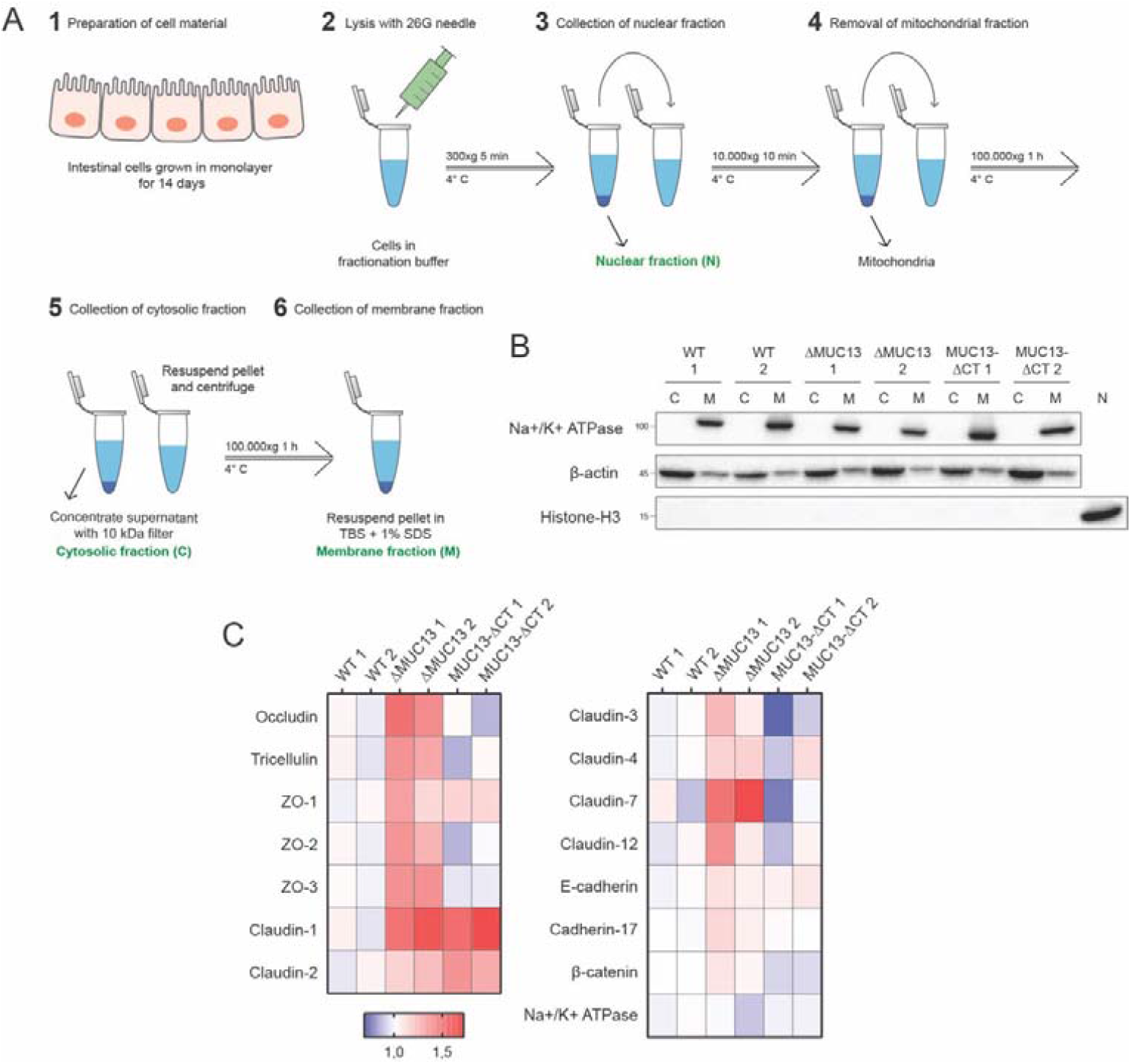
Tight junction proteins are highly upregulated in the absence of MUC13. (**A**) Subcellular fractionation protocol for the enrichment of the membrane fraction from epithelial monolayers. 1) Intestinal epithelial cell lines were grown for 2 weeks in 10 cm^2^ culture dishes. 2) Monolayers were lysed by passing through a needle in hyperosmotic fractionation buffer. 3) Nuclei (and unbroken cells) were pelleted by centrifugation and stored as the nuclear fraction (N). 4) The supernatant was collected and centrifuged again to pellet mitochondria. 5) Supernatant was again collected, and membranes were pelleted by ultracentrifugation. 6) The supernatant containing the cytosolic fraction (C) was stored. The pellet was washed and resuspended in fractionation buffer and pelleted by ultracentrifugation a second time to increase purity. 6) The resulting pellet was resuspended in TSB + 1% SDS buffer and stored as the membrane fraction (M). (**B**) Immunoblot analysis of subcellular fractionation of two WT, ΔMUC13, and MUC13-ΔCT cell lines using Na^+^/K^+^-ATPase (membrane marker), Histone-H3 (nuclear marker), and β-actin (cytoplasmic marker). C (cytosolic fraction), M (membrane fraction), N (nuclear fraction). Molecular mass standards (kDa) are indicated on the left. (**C**) Relative abundance of cell junction proteins identified by mass spectrometry in membrane fractions of WT, ΔMUC13, and MUC13-ΔCT monolayers grown for 2 weeks.

Within this group of identified proteins, 1.189 proteins had at least one membrane annotation, suggesting that the ultracentrifugation significantly enriched the membrane fractions and was important in increasing the coverage of plasma membrane and plasma membrane-recruited proteins. Upon data normalization, the intensity of the marker protein Na+/K+-ATPase was consistent between samples and biological replicates, demonstrating stringent and reproducible membrane profiling across different MUC13 mutant lines. By quantitative comparison of WT, ΔMUC13, and MUC13-ΔCT membranes by mass spectrometry, we observed a striking increase in junction proteins, as depicted in Fig. 4C. The TJ proteins occludin, tricellulin, ZO-1, ZO-2, ZO-3, and several claudins (claudins-1, -2, -3, -4, -7, and -12) were found at higher levels in ΔMUC13 membranes compared to WT membranes. We also noted an upregulation of the AJ proteins E-cadherin, β-catenin, and Cadherin-17 in ΔMUC13 membranes compared to WT membranes, though this difference was less pronounced than the upregulation of TJ proteins. In the membranes of the MUC13-ΔCT cell lines, ZO-1 and claudins-1 and -2 were consistently more abundant compared to WT membranes. The identified major tight junctions’ alterations in the ΔMUC13 and MUC13-ΔCT cell lines could underly the observed restriction of the paracellular pathway upon deletion of MUC13 or removal of its cytoplasmic tail.

### The degradation rate of tight junction proteins is not affected by MUC13

TJs are dynamic complexes in which proteins can be added and removed at different rates and quantities via vesicular transport (*20*). Internalized proteins are transported to early endosomes, followed by either trafficking to recycling endosomes to end up back at the TJ, or into late endosomes for degradation (*46, 47*). To assess the turnover of TJ proteins, monolayers were incubated with sulfo-NHS SS-biotin to label all extracellularly exposed proteins, including tight junction proteins. Cells were harvested after 1 h, 1 day, and 3 days of incubation. Biotinylated proteins were isolated from whole-cell lysates with streptavidin beads and analyzed by Western blot with specific antibodies. As before, the TJ proteins were more abundant in the ΔMUC13 cells than in WT and MUC13-ΔCT cells. In WT and MUC13-ΔCT cells, biotinylated occludin was lost after 1 day while it was still detectable in ΔMUC13 cells (Fig. 5A). Also, claudin-1 and claudin-4 were detectable for a longer period in ΔMUC13 cells compared to WT. However, quantification demonstrated that the possibility to detect proteins at day 1 was caused by the higher starting concentration, since an equal degradation rate was seen for the TJ proteins occludin, claudins-1, and claudin-4, as well as of the AJ protein E-cadherin in WT, ΔMUC13, and MUC13-ΔCT cells (Fig. 5B). These data indicate that the rate of degradation of tight junction proteins is comparable between the different cell lines, and that the increased levels of junction proteins are not due to reduced degradation.

**Fig. 5.**
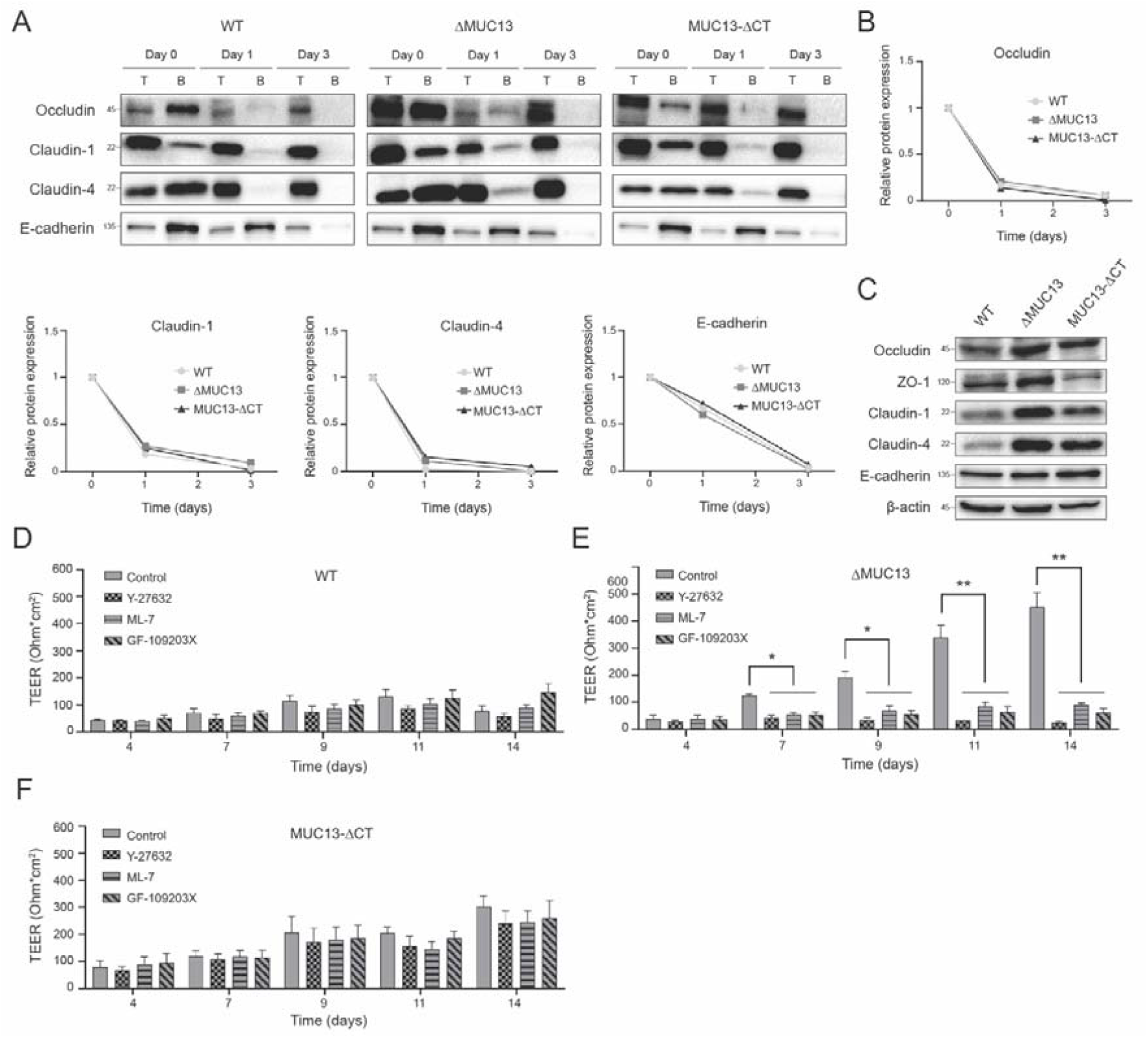
TEER buildup in the absence of MUC13 is dependent on MLCK, ROCK, and PKC kinases. (**A**) Degradation rates of biotinylated occludin, claudin-1, claudin-4, and E-cadherin analyzed by immunoblot in cell monolayers. Cells were incubated with biotin-NHS on day 0 and the presence of biotinylated proteins was determined on days 0, 1, and 3. T (total lysate), B (elution from streptavidin beads). The assay was performed at least three times and representative images are shown. Molecular mass standards (kDa) are indicated on the left. (**B**) Relative protein abundance of biotinylated occludin, claudin-1, claudin-4, and E-cadherin proteins on days 0, 1, and 3. (**C**) Immunoblot of occludin, ZO-1, claudin-1, claudin-4, E-cadherin, and β-actin in total lysates of monolayers grown for 2 weeks. The assay was performed at least three times and representative images are shown. Molecular mass standards (kDa) are indicated on the left. (**D-F**) TEER buildup of WT (D), ΔMUC13 (E), and MUC13-ΔCT (F) cell lines over time in the presence of kinase inhibitors ML-7 (MLCK), Y-27632 (ROCK), and GF-109203X (PKC). Inhibitors were added on days 3, 6, and 9 at a concentration of 50 mM (ML-7 and Y-27632) and 20 mM (GF-109203X). One representative clone for each cell line was used in these experiments. Bars represent the average and SEM of three independent experiments. *, p<0.05; ** p<0.01.

### Total tight junction protein abundance is increased in the absence of MUC13

To investigate whether these changes result from increased total expression or selective recruitment to the plasma membrane, we analyzed the abundance of selected junction proteins in the total lysates of the different cell lines by immunoblot. An upregulation of occludin, ZO-1, claudin-1, and claudin-4 was detected in ΔMUC13 lysates, whereas MUC13-ΔCT lysates displayed increased levels of occludin and claudins-1 and -4 compared to WT (Fig. 5C). Taken together, these results indicate that the increased accumulation of TJ proteins at the membrane of ΔMUC13 and MUC13-ΔCT cell lines are the result of higher total protein abundance.

### TEER buildup in the absence of MUC13 is dependent on MLCK, ROCK, and PKC kinases

The assembly, disassembly, and maintenance of TJs are known to be regulated by the kinases Myosin Light Chain Kinase (MLCK), Rho-associated protein kinase (ROCK), and members of the protein kinase C (PKC) family (*48*). MLCK, ROCK, and PKCs are all involved in the phosphorylation of MLC2, a key protein in the contraction and relaxation of the perijunctional actomyosin ring, a mechanism needed for TEER formation. In addition, PKC members phosphorylate different TJ proteins resulting in enhanced stability (*49*–*51*). For inhibition of the three kinases, we selected inhibitors ML-7 (MLCK), Y-27632 (ROCK), and GF-109293X (PKC). WT, ΔMUC13, and MUC13-ΔCT cell lines were grown for 14 days in the presence of inhibitors added on days 3, 6, and 9. The inhibitors did not have a significant effect on the TEER of WT or MUC13-ΔCT cells (Fig. 5D, F). In ΔMUC13 cells, on the other hand, the enhanced TEER buildup was not taking place in the presence of any of the three inhibitors (Fig. 5E). These results suggest that all three kinases are essential for the increased TEER buildup in the absence of the full-length MUC13.

### The effect of MUC13 on TJs is mediated by PKC proteins

We used the NetPhos-3.1 software to predict putative phosphorylation sites in the MUC13 tail and found two putative PKC binding motifs. The first motif (-VTARS-) has been previously suggested (*4*). Because this motif is situated directly adjacent to the MUC13 transmembrane domain, accessibility for PKC binding seems unlikely. The second putative PKC site -RITASRDSQ-is further removed from the transmembrane domain (amino acid residues 488-496). The truncated MUC13-ΔCT still contains the -VTARS-motif, but the -RITASRDSQ-motif is absent due to the cytoplasmic tail deletion (Fig. 6A).

**Fig. 6.**
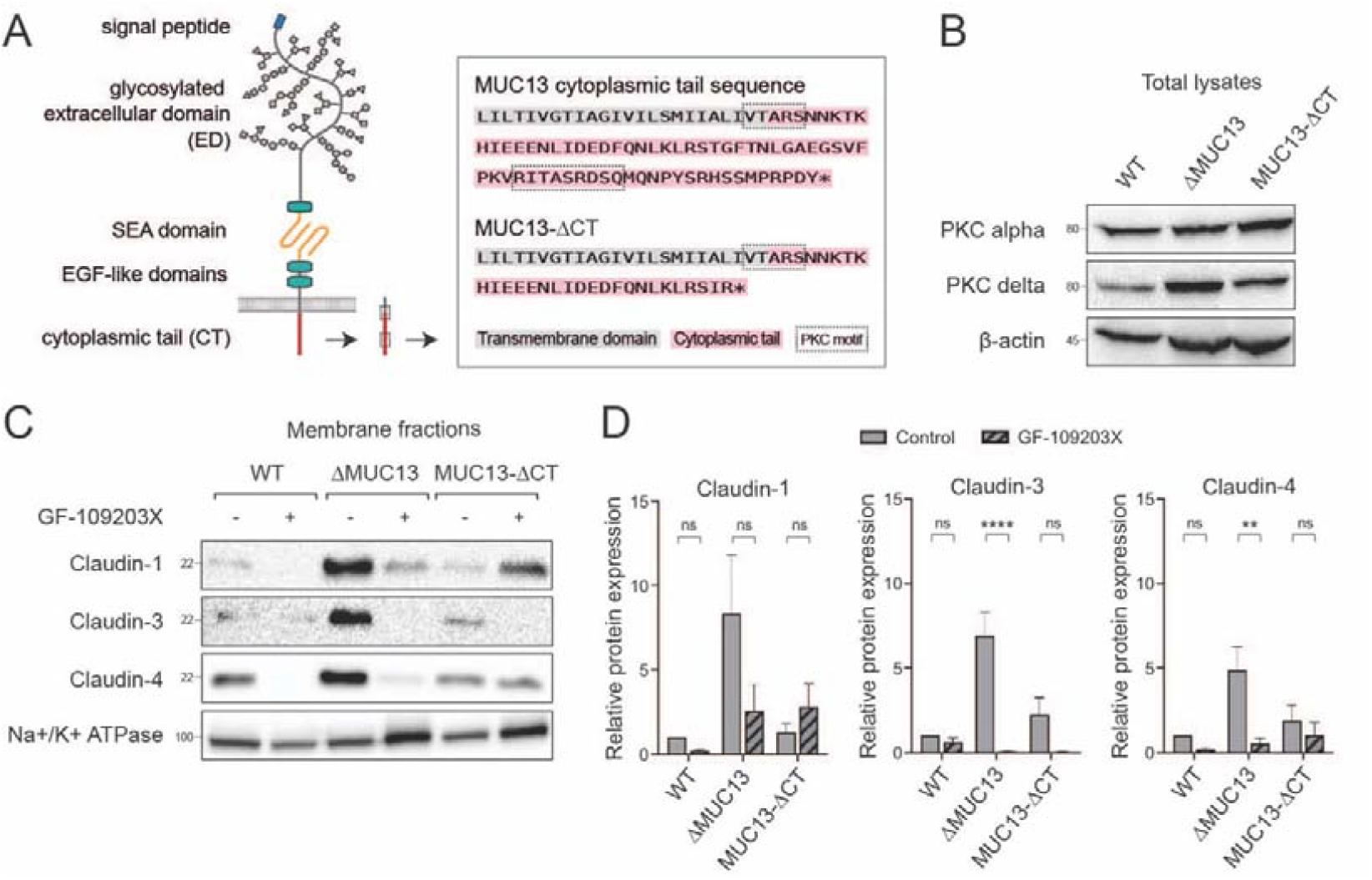
PKCs are involved in TJ regulation in the absence of MUC13. (**A**) Schematic representation of WT MUC13 domain structure (left) and protein sequence (right). The transmembrane domain (grey), the cytoplasmic tail (red), and two predicted PKC binding motifs (black boxes) are marked. (B) Immunoblot analysis of PKCα, PKCδ, and β-actin in total lysates of monolayers grown for 2 weeks. Molecular mass standards (kDa) are indicated on the left. (C) Immunoblot analysis of isolated membrane fractions from monolayers grown for 2 weeks in the presence/absence of 20 mM PKC inhibitor (GF-109203X) added every 3 days. Claudin-1, claudin-3, claudin-4, and the control protein Na^+^/K^+^-ATPase are shown. Molecular mass standards (kDa) are indicated on the left. (D) Quantification of relative protein expression of claudin-1, claudin-3, and claudin-4 in isolated fractions of cells grown in the presence/absence of GF-109203X as depicted in C. All assays were performed at least three times and representative images are shown. One representative clone for each cell line was used in these experiments. Bars represent average and SEM of three independent experiments. ns, non-significant; ** p<0.01; *** p<0.001.

To further investigate the contribution of PKC to the MUC13-related TJ phenotype, we investigated the protein expression levels of PKC isotypes PKCα and PKCδ. PKCα expression was comparable between the cell lines, but PKCδ levels were increased in the ΔMUC13 cell line (Fig. 6B). These data suggest a functional link between MUC13 and PKCδ but do not conclusively establish a connection with the PKC motif in the cytoplasmic tail. We next collected membrane fractions from WT, ΔMUC13, and MUC13-ΔCT monolayers differentiated in the absence and presence of the PKC inhibitor GF-109293X. PKC inhibition clearly resulted in reduced expression of barrier-forming claudin-1, claudin-3, and claudin-4 in the ΔMUC13 membrane fractions, and some reductions were observed in WT and MUC13-ΔCT cells (Fig. 6C). Quantification of claudin-1, claudin-3, and claudin-4 in three independent experiments demonstrated a significant reduction of claudin-3 and claudin-4 in membrane fractions of the ΔMUC13 monolayers, while the other differences were not significantly reduced. These data demonstrate that deletion of MUC13 promotes TEER buildup through increased synthesis and accumulation of TJ proteins in a PKC-dependent manner.

## Discussion

MUC13 is one of the most ubiquitously expressed transmembrane mucins in the intestinal tract, but the role of MUC13 in intestinal health and disease is not fully understood. This study explores the contribution of MUC13 to the development of intestinal epithelial barrier integrity. We provide evidence that MUC13 negatively regulates the assembly of TJ complexes and regulates the paracellular transport of small solutes. The increase in TEER observed in MUC13 knockout cells requires the signaling molecule PKC.

One of the main functions of mucins is to protect the mucosal epithelium and underlying tissues against luminal agents. In addition, the modulation of cell-cell interactions seems to be a general trait of transmembrane mucins. Several TM mucins have been implicated in the regulation of AJ proteins E-cadherin and β-catenin, namely MUC1 (*29, 34*), MUC4 (*31, 32*), MUC13 (*36*), and MUC16 (*30*). Additionally, mucin knockdown studies demonstrated the roles of several TM mucins in the regulation of TJ proteins. Silencing of MUC1 in human bronchial epithelial cells BEAS-2B led to reduced levels of occludin and claudin-1 (*37*). MUC16 knockdown in human corneal cells resulted in disruption of ZO-1 and occludin proteins, decreased TEER, and increased dye and bacterial penetration (*38, 39*). MUC17 silencing in Caco-2 and HT29-19A cells resulted in a profound reduction of occludin and ZO-1 levels and an increase in paracellular permeability after infection with enteroinvasive *Escherichia coli* (EIEC) when compared to wild type cells (*40*). In contrast to MUC1, MUC16, and MUC17 which positively regulate tight junction proteins, our study shows, for the first time, a transmembrane mucin (MUC13) that negatively regulates TJs and epithelial barrier integrity.

One of the most important functions of the intestinal epithelium is to transport nutrients and water to the mucosal tissues, while preventing the diffusion of toxins, allergens, and inflammatory molecules, such as LPS (*52*). The overall tightness or leakiness of a cell layer depends on the tight junction composition within the membrane (*53, 54*). We discovered that the removal of MUC13 causes an increased accumulation of claudins at the cell membrane, including claudins-1, -2, -3, -4, - 7, and -12 (Fig. 4C). This group contains one pore-forming claudin (claudin-2), one claudin with yet unknown barrier effect (claudin-12), and is dominated by the barrier-forming claudins-1, -3, -4, and - 7 (*20*), together resulting in the phenotypic buildup of TEER in ΔMUC13 cells compared to WT cells (Fig. 3B, C). Deletion of just the MUC13 cytoplasmic tail was sufficient to increase the accumulation of claudins-1, -2, -3, and -4. Besides ions and water, TJ proteins can regulate the paracellular flux of bigger particles through the “leak pathway”. Occludin and tricellulin have been shown to regulate the transepithelial flux of particles of various sizes (*55*–*57*). In our study, all three tested intestinal cell lines were highly restrictive for the passage of large particles, including LPS and FITC-dextran of 4 kDa and 70 kDa, but WT cells were permeable to the 520 Da Lucifer Yellow tracer while MUC13 knockout cells were not (Fig. 3G, H). Together, our results show that, in the absence of a fully functional MUC13, there is a reduction in the passage of ions and small-sized particles to deeper layers of the intestinal tissue. Alterations in ion fluxes through the paracellular channels have been described to lead to a dysfunctional intestinal barrier, causing diarrhea and malabsorption of nutrients (*58, 59*). Our observations about the link between MUC13 and TJs shed a new light on previous reports showing alterations of TJ proteins in IBD patients. The main TJ proteins occludin and tricellulin, together with sealing claudins (such as claudins-3, -4, -5, -7, and -8) are downregulated in colonic and rectal tissue of IBD patients, while claudin-1 and the pore-forming claudin-2 are upregulated leading to reduced barrier function (*28, 60*–*63*). The negative regulation of intestinal barrier MUC13 that we observe in our model may in part explain the loss of intestinal integrity in IBD patients.

The effect of MUC13 deletion on claudin expression at the membrane does not depend on altered turnover of TJ proteins (Fig. 5A-B), but rather requires kinases known to be involved in the buildup of TJs. We found that MLCK and ROCK are necessary to build up the TJ complexes and TEER in ΔMUC13 cells (Fig. 5E). These kinases are known to control the contraction of the perijunctional actomyosin ring and subsequent paracellular permeability (*48, 64*). Moreover, members of the PKC family have been implicated in the regulation of many different TJ proteins as they are responsible for the phosphorylation of claudins (Franke et al., 2010; Yoo et al., 2003; Banan et al., 2005; González-Mariscal et al., 2008, 2010), occludin (*51, 68, 69*), and ZO-1 (*49, 70*). Our study demonstrates that PKC activity is needed to accumulate high levels of claudins at the membrane of all three cell lines but is especially important in ΔMUC13 cells (Fig. 6C-D). The MUC13 cytoplasmic tail contains several potential phosphorylation sites (8 serine and 2 tyrosine residues) (*2, 4*) and two putative PKC motifs (Fig. 6A). We observed that in the absence of the full-length MUC13, the total levels of PKCδ are increased but deletion of the cytoplasmic tail alone did not evidently increase PKCδ levels (Fig. 6B). On the other hand, PKC inhibition did reduce the expression of some TJ proteins in the MUC13-ΔCT cells, but these changes were not significant between three experiments (Fig. 6C, D). This might suggest that deletion of the MUC13 tail only can enhance PKC activity to stabilize TJ proteins. A role for PKCδ in the regulation of TJs is in line with previously reported results linking PKCδ to upregulation of claudin-1 and claudin-7 protein levels in different epithelial cell lines (*51, 71, 72*). In future studies, we intend to address the link between MUC13 and PKCδ during TJ regulation in more detail. Based on our current data, we propose a model in which MUC13 negatively impacts claudin build up at the membrane by regulating the levels and/or activity of PKCδ (Fig. 7A-C).

**Fig. 7.**
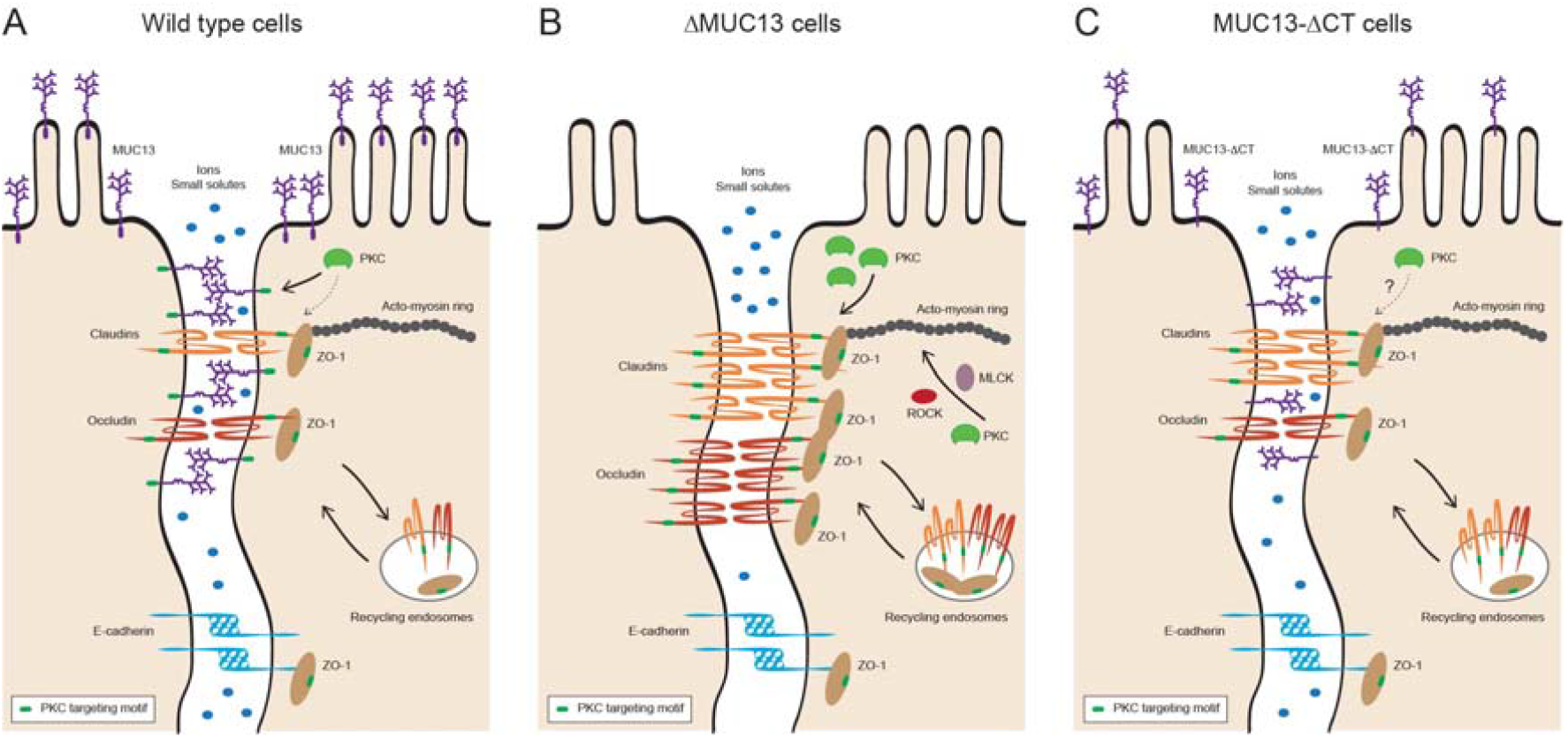
Tight junction regulation by MUC13. (**A**) In wild type cells, MUC13 localizes to both the apical surface and tight junction (TJ) region of the lateral membrane. Cell junction complexes that contain claudins, occludin, ZOs, and E-cadherin, are assembled along the lateral membrane. Under normal conditions, there is some paracellular passage of ions and small solutes, a process that is controlled by the TJ proteins claudins and occludin. MUC13 cytoplasmic tail has a putative PKC binding motif, which may play are role in recruiting PKC and controlling its activity and/or stability. Cell junction proteins such as claudins, occludin, and ZO-1 also can be targeted by PKCs. (**B**) In the absence of the complete MUC13 protein, TJ proteins (occludin, claudins, and ZO-1) are accumulating at the membrane over time, causing increased transepithelial resistance (TEER) and lower paracellular passage of small solutes. The TEER buildup in ΔMUC13 cells is dependent on MLCK, ROCK and PKC kinases. The accumulation of claudins at the membrane in ΔMUC13 cells is PKC-dependent and is not caused by slower degradation rates of TJ proteins through recycling endosomes. (**C**) Removal of the MUC13 cytoplasmic tail leads to an intermediate phenotype with some accumulation of claudin-1, -3, -4, and ZO-1 at the membrane, but to a lower extent compared to the full knockout. The role of PKC in this cell line remains to be determined. MUC13-ΔCT cells are less permeable to small solutes but do not show a significant increase in TEER when compared to WT cells. The degradation rate of TJ proteins in MUC13-ΔCT cells is comparable to WT and ΔMUC13.

Confirmation of the MUC13 cytoplasmic tail deletion cell lines with MUC13-targeted antibodies indicated that both MUC13-ΔCT cell lines lacked the cytoplasmic tail. The MUC13 extracellular domain was detectable in the MUC13-ΔCT 1 cell line but expression in the MUC13-ΔCT 2 cell line was very limited (Fig. 2F, G). The reduced percentage of MUC13-ΔCT cells expressing MUC13 on the surface compared to WT cells might indicate reduced stability of the MUC13 protein lacking the cytoplasmic tail. Despite the possible differences in MUC13 stability or antibody binding, the phenotypic characterization of both MUC13-ΔCT cells was very similar and neither cell line phenocopied the effect of the full MUC13 knockout on TEER and TJ. Therefore, we are confident that the targeted removal of the MUC13 cytoplasmic tail was successful, and the results obtained with these cell lines are reliable. Removal of the MUC13 cytoplasmic tail led to a partial phenotype, where ZO-1 and claudins-1, -2, -3, and -4 were upregulated in the membrane, although to a lower extent compared to the full knockout. This was accompanied by a slight increase in TEER and reduced paracellular permeability to Lucifer Yellow substrate compared to WT cells. Also in the PKC experiments, the MUC13-ΔCT cells showed a partial phenotype but did demonstrate a dependency on PKC activity for elevated expression of TJ proteins. Together, these results point to a pivotal role for both the MUC13 extracellular domain and cytoplasmic tail in TJ buildup and underline the challenge of studying the functions of different TM mucin domains.

In healthy conditions, the transmembrane mucin MUC13 is involved in important biological processes, including cell growth and maintenance (*16, 17*), protection of cells from toxin-induced damage (*11*), and formation of a physical barrier to reduce bacterial contact (*2*). MUC13 is upregulated during IBD (*11, 73*) and CRC (*10, 74*), correlating with increased pro-inflammatory responses (*75*), cell growth, and migration (*16, 17*). Our results demonstrate that MUC13 is a negative regulator of PKC-mediated TJ protein assembly at the membrane. Overexpression of MUC13, as observed in IBD, may lead to a reduction in TJ proteins, such as occludin, claudins, and ZOs, and increased paracellular permeability to water, ions, and organic solutes. Opening of TJ complexes is essential to allow sampling of luminal bacteria by immune cells but decreased barrier function can also contribute to the development of chronic intestinal inflammation. It is interesting to speculate that MUC13 with its complex extracellular domain, could play a role in sensing the inflammatory state of the intestine and can respond by regulating TJs through its cytoplasmic tail. Our study brings to light that the transmembrane mucin MUC13 plays a unique role in the intestinal epithelium and emphasizes the need for further studies into the functions of specific mucins.

## Materials and Methods

### Cell lines, bacteria, and culture conditions

The human intestinal epithelial cell lines HRT18 (ATCC-CCL-244), Caco-2 (ATCC-HTB-37), and CRISPR/Cas9 knockout derivative cell lines used in this study, as well as HEK293T cells (CRL-3216, ATCC) were routinely grown in 25 cm^2^ flasks in Dulbecco’s modified Eagle’s medium (DMEM) + glutamax (Life technologies, 31966047) containing 10% fetal calf serum (FCS) (Sigma, F7524) at 37°C in 10% CO_2_. HEK-Blue™ Null and HEK-Blue™ hTLR4 cells were purchased from InvivoGen (hkb-htlr4) and cultured in DMEM containing 10% heat-inactivated FCS, penicillin/streptomycin (BioConnect, ML-105L), and antibiotics from Invivogen (Zeozin (ant-zn) and Normocin (ant-nr) for HEK-Blue™ Null cells; Zeozin, Normocin, Blasticidin (ant-bl), and Hygromycin (ant-hg) for HEK-Blue™ hTLR4 at 37°C in 5% CO_2_. Cells were detached with 0.25% trypsin (ThermoFisher, 25200-072), passaged twice a week in a 1:10 dilution and split before they reached 80% confluency. *Lactobacillus plantarum* (ATCC, 14917) was grown in MRS medium in aerobic conditions.

### Antibodies and reagents

For Western blotting, antibodies against claudin-1 (ThermoFisher, 51-9000), claudin-3 (ThermoFisher, 34-1700), claudin-4 (ThermoFisher, 32-9400), occludin (Invitrogen, 33-1500), Zonula Occludens-1 (ZO-1) (Abcam, ab216880), E-cadherin (Abcam, ab1416), PKCα (ab32376), PKCδ (ab182126), b-actin (Bioss, bs-0061R), MUC13 (Abcam, ab235450), MUC13 hybridoma supernatant (in house), Na^+^/K^+^-ATPase (Abcam, ab76020), and acetyl-Histone H3K9 (Merck, 07-352) were used. Secondary antibodies used for immunoblotting were goat anti-mouse-HRP (Sigma, A2304) and goat anti-rabbit-HRP (Sigma, A4914). Secondary antibodies used for immunofluorescence were goat anti-mouse-Alexa488 (ThermoFisher, A11029), goat anti-mouse-Alexa568 (ThermoFisher, A11031), goat anti-rabbit-Alexa488 (ThermoFisher, A11034), goat anti-mouse-Alexa568 (ThermoFisher, A11036), and DAPI (D21490, Invitrogen).

For permeability assays, 4 kDa (46944) and 70 kDa (46945) Fluorescein isothiocyanate-dextran (FITC), and Lucifer Yellow CH dipotassium salt (L0144) were purchased from Sigma. Ultrapure LPS from *Escherichia coli* was purchased from InvivoGen (tlrl-3pelps). For biotinylation assays, Pierce™ Premium Grade sulfo-NHS SS-biotin was acquired from ThermoFisher (PG82077).

### Bioinformatics Single Cell Studies

Single cell gene expression from intestinal epithelial cells was analyzed using a public single cell RNA-sequencing dataset (*76*). The H5AD file containing data from all epithelial cells was downloaded from https://www.gutcellatlas.org and further analyzed in Rstudio using the packages “Seurat”, “SeuratData”, and “SeuratDisk”. Cells from healthy adult subjects were selected, and low-quality cells (less than 2000 genes or >20% of counts mapping to mitochondrial genes) were removed. Data from the remaining 37,325 cells were then normalized using the SCTransform algorithm and dotplots showing the expression by cell type or by intestinal zone were made. Rare cell types (less than 100 in the dataset) are not shown in the plots.

### Generation of HRT18 ΔMUC13 and MUC13-ΔCT cell lines using CRISPR/Cas9

To generate ΔMUC13 cells, we used the pCRISPR-hCas9-2xgRNA-Puro plasmid (Langereis *et al*., 2015) that encodes Cas9 with two MUC13-specific guide RNAs to generate a 380 bp deletion in the second exon of the *MUC13* gene. The pCRISPR plasmid was digested with *Sap*I and simultaneously dephosphorylated with alkaline phosphatase (FastAP; ThermoFisher). Guide RNA primer sets A (KS40 5’-ACCGACCACAGAAACTGCGACTAG -3’ and KS41 5’-AACCTAGTCGCAGTTTCTGTGGTC-3’) and B (KS42 5’-CCGTCCCACTGGCACCGCTTTATG-3’ and KS43 5’-AAACATAAAGCGGTGCCAGTGGGA-3’) were phosphorylated with T4 polynucleotide kinase (ThermoFisher) at 37°C for 30 min and annealed by cooling down from 85°C to 25°C at 0.1°C/sec. Annealed primer sets were ligated into the *Sap*I-digested pCRISPR plasmid and confirmed by sequencing with primers KS46 5’-GTTCACGTAGTGCCAAGGTCG-3’ and KS47 5’-GAGTCAGTGAGCGAGGAAGC-3’, resulting in plasmid pCR4. Two-day grown HRT18 cells were trypsinized from a 25 cm^2^ flask and transfected in suspension with 2 μg of pCR4, pCRISPR-empty, or no plasmid using Fugene (Promega) according to the manufacturer’s instructions. Cells were cultured in DMEM + 10% FCS for two days, after which 5 μg/ml puromycin (Life Technologies) was added to the medium to select for positively transfected cells. Cells were maintained in medium with puromycin until all negative control cells had died. Single-cell cloning was performed by serial dilution and single-cell clones were tested for the MUC13 deletion by PCR with primers KS126 5’-CCAGGGGTTTATGACCAATCTAGG-3’ and KS127 5’-TGCACAGCTAGCAAATAACTTGAGG-3’. The deletion in the MUC13 was confirmed by sequencing and the clones were named HRT18-ΔMUC13 clones 1 and 2. The cells transfected with the empty CRISPR plasmid served as control in all experiments and were renamed wild type (WT) for clarity of the figures. The absence of MUC13 protein in the knockout cell lines was confirmed by immunoblot with anti-MUC13 antibody. To generate the MUC13-ΔCT cell line, a similar protocol as described above was followed with guide RNA primer sets A (CSP5 5’-ACCGAATCTAAAACTGCGGTCGAC -3’ and CSP6 5’-AACGTCGACCGCAGTTTTAGATTCC -3’) and B (CSP7 5’-CCGGCACTGACTCACCTAATAGTCG -3’ and CSP8 5’-AAACGACTATTAGGTGAGTCAGTGC -3’) to generate a deletion of 121 bp in the tenth exon of the *MUC13* gene. The resulting single clones after transfection were confirmed with primers CSP96 5’-TCAAGTGATCTGCCCACCACGG-3’ and CSP97 5’-TCTGCCCTGGTGCATTCACTCC-3’.

### Overexpression of MUC13 in HRT18-WT and HRT18-ΔMUC13 cells

Cloning of the original MUC13 gene sequence in *E. coli* DH5a was problematic. The Softberry promoter prediction algorithm (*77*) was used to analyze the MUC13 sequence which revealed a multitude of predicted Sigma70 binding sites. We altered the MUC13 sequence with synonymous mutations to remove the predicted binding sites. The optimized MUC13 sequence was ordered from Thermo Fisher. To generate doxycycline-inducible expression of the MUC13-opt constructs, the plasmid pInducer 20-extended MCS (pKSU59) (*78*) was used as a vector. First, restriction sites were added via PCR amplification with primers XL14-Fwd (5’ CCGCTCGAGGCCACCATGGAAGCCATCATTCATCTTACTCTTC 3’) and XL13-Rev (5’ TATGGCGCGCCCCATAGAGCCCACCGCATC 3’) to obtain the insert fragment (MUC13opt-GFP) from plasmid pDS2 (pcDNA3.1-MUC13opt-GFP). Then, both insert and vector were digested with Ascl-FD and Xhol-FD ligated together to generate plasmid pJSU002 (pInducer20-MUC13opt-GFP). This plasmid was subsequently used to generate inducible overexpression of MUC13 in HRT18-WT and HRT18-ΔMUC13 cells using lentiviral transduction. For lentiviral production, HEK293T cells were seeded at 70% confluence in 6-well tissue culture plates 24 h before transfection. Lipofectamine ™ 3000 (Invitrogen, L3000001) was used as the transfection reagent according to the manufacture’s protocol. Cells were incubated with the transfection mix for 6 h, media was replaced with fresh DMEM/10% FCS and grown for 48 h. The subsequent steps for lentivirus transduction on HRT18-WT and HRT18-ΔMUC13 cells were performed as previously described (*78*). The resulting cells were called HRT18-WT+pMUC13 and HRT18-ΔMUC13+pMUC13 cells. Cells were also transfected with empty pInducer plasmid as controls, resulting in HRT18-WT Ctr and HRT18-ΔMUC13 Ctr cell lines. To validate the expression of MUC13opt-GFP constructs, cells were induced with 20 ng/ml of doxycycline (Sigma, D3072) for 24 h and observed under a fluorescent microscope for GFP signal.

### Immunofluorescence and confocal microscopy

For immunofluorescence, cells were grown on coverslips in 24-well plates for 14 days. Monolayers were washed twice with Dulbecco’s Phosphate Buffered Saline (DPBS, Sigma, D8537) and fixed with 4% cold paraformaldehyde in PBS (VWR, J19943) for 30 min at room temperature (RT). The fixation was stopped by incubation with 50 mM NH_4_Cl in PBS for 10 minutes. Cells were washed twice with DPBS before they were incubated with primary antibodies (MUC13 at 1:100 dilution, occludin at 1:50, and E-cadherin at 1:100) in binding buffer (0.2% Triton-X100 (Sigma, X100), 2.2% gelatin (Sigma, CM135a), and 0.2% BSA (Sigma, A7030) in DPBS for 1h at RT. Coverslips were washed 3 times with binding buffer followed by incubation with secondary antibodies (1:200) and DAPI (1:500) for 1h at RT. Coverslips were washed 3 times with DPBS, once with MilliQ, and embedded in Prolong diamond mounting solution (ThermoFisher, P36961). Images were collected on a Leica SPE-II confocal microscope in combination with Leica LAS AF software. Image analysis was performed using Fiji/ImageJ.

### Transepithelial electrical resistance (TEER) measurements

Cells were seeded in 12-well Transwell plates with 12 mm inserts and 0.4 mM membrane pore size (Costar, 3401) at 30%, 40%, and 60% confluency, respectively. Wells without cells were taken along as negative control. Transepithelial electrical resistance was determined with a Millicell ERS-2 Voltohmmeter (Millipore). All measurements were performed on three individual wells. TEER measurements were taken every 2-3 days for 2 weeks. TEER W/cm^2^ values were calculated by subtracting the average negative control value from the measurement and multiplying it by the well surface (1,12 cm^2^). For the MUC13 overexpression experiments, HRT18-WT Ctr, HRT18-WT+pMUC13, HRT18-ΔMUC13 Ctr, and HRT18-ΔMUC13+pMUC13 cells were seeded in 24 Transwell plates with 6.5 mm inserts and 0.4 mM membrane pore size (Costar, 3470) at 30% (WT) and 60% (ΔMUC13) confluency. TEER was measured every 2-3 days for 2 weeks. At day 14, doxycycline was added to the top compartment at a concentration of 0.2 μg/mL. To study the effect of MLCK, ROCK, and PKC on TEER build up over time, ML-7 (50 mM), Y-27632 (50 mM), or GF-109203X (20 mM) inhibitors were added to the upper compartment, respectively, at days 3, 6, and 9. TEER measurements were taken every 1-2 days for 2 weeks.

### Epithelial permeability assays with Lucifer Yellow, FITC-Dextran and LPS

Epithelial paracellular permeability for particles was assessed by measuring the flux of 0.5 kDa Lucifer Yellow CH dipotassium salt and 4 and 70 kDa FITC-Dextran particles across confluent monolayers. Cells were grown for 2 weeks in 12-well Transwell plates with 12 mm inserts and 0.4 mM membrane pore size (Costar, 3401). To minimize interference from the media when measuring FITC, media from the bottom wells was changed to DMEM without red phenol + 10% FCS. Subsequently, 500 mL of 4 or 70 kDa FITC-Dextran dissolved to 1 mg/mL in DMEM without red phenol or 500 mL of 400 mg/mL LY was added to the top compartments. After 2 h incubation with LY or 6 h with FITC-Dextran particles, 100 mL aliquots were taken from the bottom wells and the fluorescent intensity was measured with a FLUOstar Omega Microplate Reader (BMG Labtech). The excitation and emission wavelengths were 492 nm and 520 nm for FITC-Dextran, and 428 and 540 nm for LY. The percentage of permeability was calculated by comparing the fluorescence intensity to that of membrane-only wells.

### LPS translocation assays

Cells were grown for 2 weeks in 12-well Transwell plates with 12 mm inserts and 0.4 mM membrane pore size (Costar, 3401). The media from the bottom compartment was changed to 500 mL DMEM without FCS. 5 mg of Ultrapure *E. coli* LPS diluted in DMEM without FCS was added to the top wells and incubated at 37°C for 24 h. To determine the maximum amount of LPS that could be translocated, 5 mg of LPS was added to wells without cells (membrane only). The next day, the bottom compartments were frozen at -20°C until further use. For quantitative detection of LPS, HEK-Blue™ hTLR4 were used with HEK-Blue™ Null cells as negative control. 2.5×10^4^ HEK-Blue™ Null cells and 3.5×10^4^ HEK-Blue™ hTLR4 cells (due to slightly slower growth of the Null cells) were seeded in 96-well flat-bottomed tissue culture plates and incubated at 37°C for 24 h. Then, cells were stimulated with 100 mL of media from the Transwell bottom of the LPS translocation experiment. For quantification of the LPS concentration, 10-fold dilutions of LPS from 100 ng/mL to 0.1 ng/mL in 100 mL were used. HEK-Blue™ hTLR4 and HEK-Blue™ Null cells were stimulated with the LPS-containing fractions for 24 h at 37°C. Relative NF-κB activity as a result of TLR4 stimulation was determined by quantifying the secreted alkaline phosphatase (SEAP) activity. Twenty mL of HEK-Blue™ supernatants were transferred to a 96-well plate containing 180 mL pre-warmed (37°C) QUANTI-Blue™ (the substrate for SEAP, InvivoGen, rep-qbs). Reactions were developed at 37°C for 50-90 min and measured at 630 nm using FLUOstar Omega Microplate Reader (BMG Lactech). Three wells with DMEM only were used as blanks and subtracted from the other measurements.

### Epithelial barrier experiments with *Lactobacillus plantarum*

Cells were grown for 2 weeks in 12-well Transwell plates with 12 mm inserts and 0.4 mM membrane pore size (Costar, 3401). An overnight culture of *L. plantarum* was added at MOI 50 at the apical side in a final volume of 500 mL in DMEM without FCS. Media in the basolateral compartment was replaced with fresh DMEM without FCS. TEER was measured at multiple time points until 42 h. All measurements were performed on three individual wells and in three independent biological replicates. TEER W/cm^2^ values were calculated by subtracting the average negative control value from the measurement and multiplying it by the well surface (1.12 cm^2^).

### Subcellular fractionation

For subcellular fractionation of epithelial monolayers, a protocol from Abcam (https://www.abcam.com/protocols/subcellular-fractionation-protocol) was used with some modifications. Cells were grown in 10 cm^2^ culture dishes for 2 weeks, washed twice with ice-cold PBS and scrapped with a cell scraper in 500 mL fractionation buffer (20 mM HEPES pH 7.4, 10 mM KCl, 2 mM MgCl_2_, 1 mM EDTA, 1 mM EGTA, 1 mM DTT, and 1x protease and phosphatase inhibitors) and transferred to an Eppendorf tube. Cell suspensions were passed 10 times through a 26G needle and centrifuged at 300 x g for 5 min. The supernatant containing cytoplasm, membranes and mitochondria was transferred to a new Eppendorf tube and kept on ice. To maximize cell membrane rupture, these steps were repeated: resuspension of the pellet in buffer, lysis by a needle, and centrifugation. The recovered supernatants were centrifuged at 10,000 x g for 10 min to separate the mitochondria (pellet) from the cytoplasm and membranes (supernatant). Supernatants were transferred to 1.5 mL microcentrifuge tubes (Beckman Coulter) and centrifuged in an ultracentrifuge at 100,000 x g for 1 h at 4°C. Supernatants containing the cytosolic fraction were transferred to a Spin-X UF 10 kDa Centrifugal Concentrator (Corning) and concentrated by centrifugation to a final volume of 100 mL. The pellet of the ultracentrifugation step containing the membrane fraction was taken up in 500 mL of fractionation buffer and re-centrifuged at 100,000 x g for 1 h at 4°C for increased purity. The pellet was resuspended in 100 mL TBS (50 mM Tris, 150 mM NaCl, 1% SDS). Protein concentrations in all fractions were determined with Pierce™ BCA Protein Assay kit (ThermoFisher, 23225) and equal amounts were loaded into SDS-PAGE gels to confirm the efficiency of the protocol. Na^+^/K^+^-ATPase protein was chosen as a control for the membrane fraction, b-actin for the cytosolic fraction, and acetyl-Histone H3K9 to exclude nuclear contamination.

### Immunoblotting

Cell pellets were taken up in 1% SDS in PBS and lysed by mechanical lysis through a 26G needle. Protein concentration was determined using a Pierce™ BCA Protein Assay kit and equal amounts of protein were prepared in Laemmli sample buffer and boiled for 5 minutes at 96°C. For immunoblotting of MUC13, protein lysates were loaded onto 8% SDS-PAGE gel and transferred to a PVDF membrane using a semi-dry transfer system (Biorad) for 10 min at 2.5 amperes. The membranes were blocked with 5% skimmed milk powder in PBS-Tween for 1 h at RT. Subsequently, the membranes were incubated with MUC13 antibody directed against CT domain (Abcam) at 1:250 dilution in PBS-Tween containing 5% skimmed milk powder o/n at 4°C. The next day, the membranes were washed 4 times with PBS-Tween (10 minutes each) and incubated with secondary antibody diluted 1:5,000 in PBS-Tween containing 5% skimmed milk powder for 1 h at RT. For immunoblotting of other proteins, protein lysates were loaded onto 8-12% SDS-PAGE gels and transferred to PVDF membranes. Blocking was done o/n at 4°C in 5% BSA-TSMT (20 mM Tris, 150 mM NaCl, 1 mM CaCl_2_, 2 mM MgCl_2_ adjusted to pH 7 with HCl and 0.1% Tween 20). Antibodies were diluted in 1% BSA-TSMT and incubated for 1 h at RT. Antibodies were used at 1:1,000 dilution, except for claudin antibodies which were used at 1:500 dilution and b-actin antibody at 1:2,000. For visualization, blots were incubated with Clarity Western ECL solution (Biorad) and imaged in a Bio-Rad Gel-Doc system.

### Cell-surface biotinylation to determine recycling of TJ proteins

Cells were grown for 10 days in 6-well Transwell plates with 24 mm inserts and 0.4 mM membrane pore size (Costar, 3412). 1 mg/mL of sulfo-NHS SS-biotin dissolved in PBS was added to the upper and basal compartments and incubated for 1 h at 4°C. Free biotin was washed away twice with cold sulfo-NHS SS-biotin blocking solution (50 mM NH4Cl in PBS, 1 mM MgCl2, 0.1 mM CaCl2). Five hunderd mL Lysis Buffer (50 mM Tris-HCl pH 7.5, 150 mM NaCl, 1% SDS, 1 mM PMSF, and EDTA-free protease inhibitor cocktail from Roche, dissolved in PBS) was added and cells were harvested from the Transwell membrane using a disposable cell scrapper and transferred to an Eppendorf tube. Samples were lysed for 45 min at RT by mechanical lysis. These samples were labeled as Day 0 (maximum amount of labeled proteins). Fresh DMEM + 10% FCS + Pen/Strep was added to the other wells and incubated at 37 °C for 1 day or 3 days, after which cells were harvested and lysed as described above. After incubation with Lysis Buffer, lysates were cleared of insoluble debris by centrifugation at 16,000 x g for 10 min. A small fraction of all cleared lysates was saved in another tube for the total protein sample. Per sample, 60 mL of Pierce Streptavidin Agarose Beads (ThermoScientific) were washed with 1 mL Lysis Buffer in a 2 mL microcentrifuge tube, and centrifuged for 2 min at 4,500 x g. After a second wash, beads were resuspended in Lysis Buffer equivalent to 60 mL/sample. Samples (20 mL) were loaded onto SDS-PAGE gels and immunoblotting was performed using claudin-1 and -4, occludin, and E-cadherin antibodies. Band intensities in each blot were analyzed with Image Lab Software 5.0.

### Sample Preparation for Mass Spectrometry

After fractionation, proteins in the membrane fraction were reduced in 10 mM dithiothreitol (DTT) at 20°C for 1 h and then alkylated with 20 mM iodoacetamide (IAA) at 20°C for 30 min in the dark. Excess IAA was quenched with an additional 10 mM DTT. Lys-C (Wako, Japan) was added at an enzyme/protein ratio of 1/75 and incubated for 4 h at 37 °C. Then, the solution was diluted with 50 mM ammonium bicarbonate to reach a 2 M final concentration of urea, and trypsin was added (Sigma, USA) at an enzyme/protein ratio of 1/75 and digested overnight at 37 °C. The digested samples were quenched with 2% formic acid on the second day and desalted with Sep-Pak C18 1 cc Vac cartridge (Waters, USA). Desalted samples were dried by vacuum centrifugation and stored at - 80 °C for further use.

### LC-MS/MS

Peptides were reconstituted in 2% formic acid and analysed in triplicates. LC-MS/MS was performed using an Orbitrap Exploris 480 mass spectrometer (Thermo Scientific) coupled with an UltiMate 3000 UHPLC system (Thermo Scientific) fitted with a μ-precolumn (C18 PepMap100, 5 μm, 100 Å, 5 mm × 300 μm; Thermo Scientific), and an analytical column (120 EC-C18, 2.7 μm, 50 cm × 75 μm; Agilent Poroshell). Peptides were loaded in solvent A (0.1% formic acid in water) with a flow rate of 30 μl/min and then separated by using a 115-min linear gradient at a flow rate of 0.3 μl/min. The gradient was as follows: 9% solvent B (0.1% formic acid in 80% acetonitrile, 20% water) for 1 min, 9– 10% in 1 min, 10–36% in 95 min, 36–99% in 3 min, 99% for 4 min, 99-9% in 1 min, and finally the system equilibrated with 9% B for 10 min. Electrospray ionization was performed by using 1.9 kV spray voltage; the temperature of the ion transfer tube was set to 275 °C, and the RF lens voltage was set to 40%. MS data were acquired in data-dependent acquisition mode. Full scan MS spectra were acquired accumulating to ‘Standard’ pre-set automated gain control (AGC) target, at a resolution of 60,000 within the m/z range of 375-1600. Multiply charged precursor ions starting from m/z 120 were selected for further fragmentation. Higher energy collision dissociation (HCD) was performed with 28% normalized collision energy (NCE), at an orbitrap resolution of 30,000. Dynamic exclusion time was set to 16 s and 1.4 m/z isolation window was used for fragmentation.

### Data analysis for MS

MaxQuant software (version 1.6.10.0) was used for raw data analysis. The database search was performed against on human UniProt database (version April 22, 2021) by using the integrated Andromeda search engine. Protein N-terminal acetylation and methionine oxidation were added as variable modifications; cysteine carbamidomethylation was added as a fixed modification. Digestion was defined as Trypsin/P and up to 2 miscleavages were allowed. Label-free quantification (LFQ) and the match-between-runs feature were applied for identification. 1% false discovery rate (FDR) was applied for both peptide and protein identification.

Quantitative data filtering was performed in the Perseus software (version 1.6.10.0). Potential contaminants and reverse peptides were removed, and all the LFQ intensities were normalized with log2 transformation. Proteins quantifiable in at least two out of three replicates were retained. Imputation was performed based on a normal distribution. A two-sided paired Student’s t-test was performed with permutation-based FDR (*q*-values) from 250 randomizations. Proteins were considered significant if *q*-values were 0.05 or less.

#### MS data availability

All the proteomics raw data were deposited to the ProteomeXchange Consortium with the dataset identifier PXD029606.

### Statistical analysis

Statistical analysis was performed in IBM SPSS Statistics version 27 and depicted by Graph Pad Prism 7 software. Kolmogorov-Smirnov test was used to assess normality of the data, and log transformation was used when the data was not normally distributed. Statistical differences in data including TEER development over time, FITC Dextran, Lucifer Yellow, and LPS translocation were analyzed using one-way ANOVA (analysis of variance) with Tukey’s HSD *post hoc* test. TEER build up in the presence of inhibitors was analyzed using two-way ANOVA with Dunnett’s *post hoc* test. The effect of *L. plantarum* on TEER was determined by calculating the fold change (42 h vs 0 h) and analyzing statistical differences using an independent t-test. All graphs depict the mean and standard error of the mean (SEM) of at least three independent experiments. Results of all performed statistical tests are depicted in the figures. A *p*-value of <0.05 was considered significant. *, p<0.05; ** p<0.01; *** p<0.001.

## Acknowledgments

We thank Dr. Richard Wubbolts and Ing. Esther van ‘t Veld from the Center for Cell Imaging of Utrecht University for assistance with microscopy.

## Author contributions

Conceptualization: CSP, DS, KS. Methodology: CSP, DS, BW, WW, KS. Investigation: CSP, DS, ZM, JS, BW, WW, KS. Visualization: CSP, KS. Supervision: KS, JVP. Writing original draft: CSP, KS. Review & editing: CSP, DS, ZM, JS, BW, WW, JVP, KS.

## Supplementary Materials

**Fig. S1.**
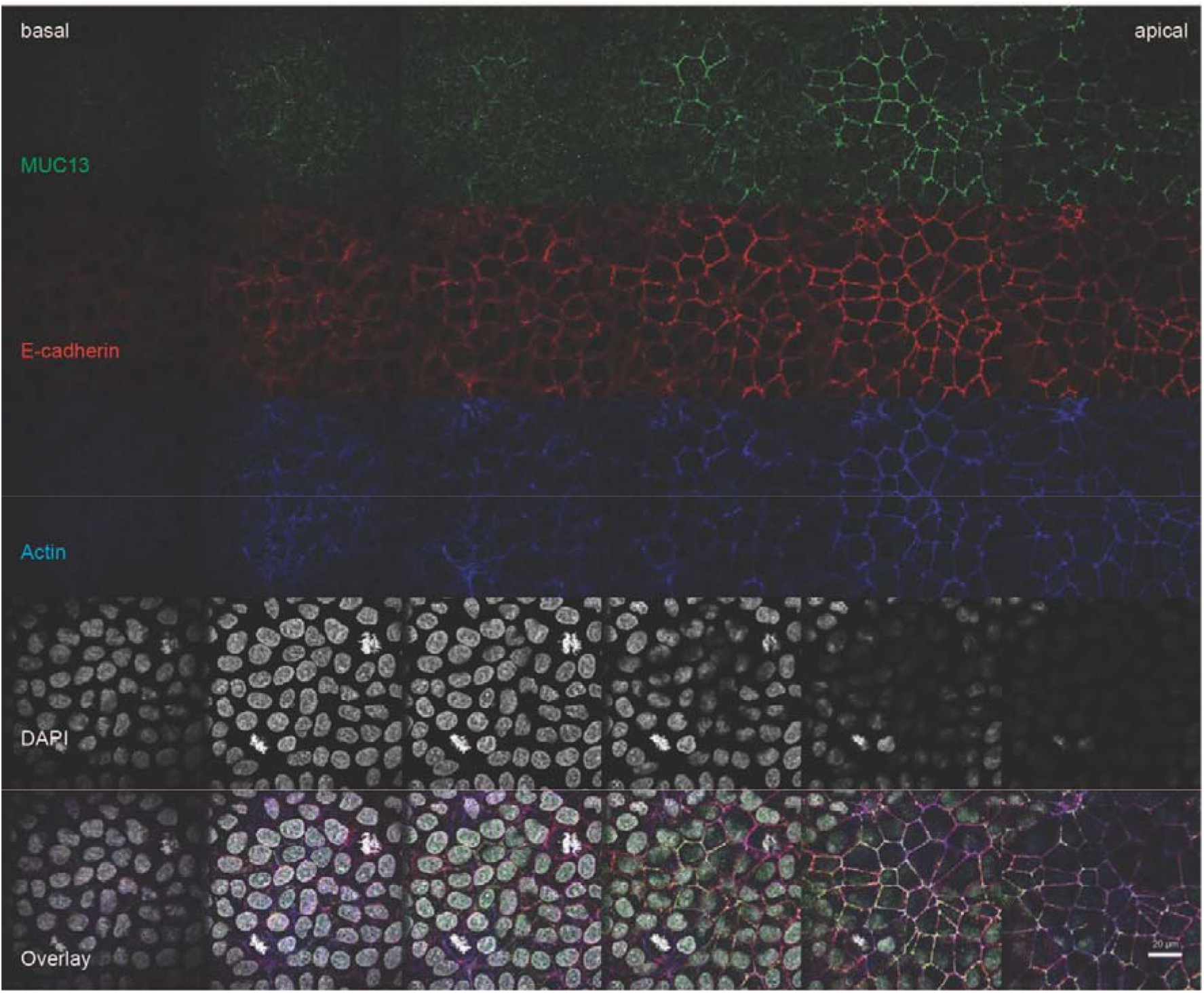
MUC13 localizes at the upper side of the lateral membrane. (**A**) Immunofluorescence of HRT18 intestinal cells stained for MUC13 cytoplasmic tail (MUC13-CT) (green), E-cadherin (red), β-actin (blue), and DAPI (white). White scale bars represent 20 mM. Pictures were taken at different heights in the epithelial monolayer (Z).

## References

1. J. J. Kim, W. I. Khan, Goblet cells and mucins: role in innate defense in enteric infections. Pathogens. 2, 55–70 (2013).

2. J. P. M. van Putten, K. Strijbis, Transmembrane Mucins: Signaling Receptors at the Intersection of Inflammation and Cancer. J Innate Immun. 9, 281–299 (2017).

3. M. E. V. Johansson, H. Sjövall, G. C. Hansson, The gastrointestinal mucus system in health and disease. Nat Rev Gastroenterol Hepatol. 10, 352–361 (2013).

4. S. J. Williams, D. H. Wreschner, M. Tran, H. J. Eyre, G. R. Sutherland, M. A. McGuckin, Muc13, a novel human cell surface mucin expressed by epithelial and hemopoietic cells. J Biol Chem. 276, 18327–18336 (2001).

5. K. L. Carraway, V. P. Ramsauer, B. Haq, C. A. Carothers Carraway, Cell signaling through membrane mucins. Bioessays. 25, 66–71 (2003).

6. P. K. Singh, M. A. Hollingsworth, Cell surface-associated mucins in signal transduction. Trends Cell Biol. 16, 467–476 (2006).

7. M. Gersemann, S. Becker, I. Kübler, M. Koslowski, G. Wang, K. R. Herrlinger, J. Griger, P. Fritz, K. Fellermann, M. Schwab, J. Wehkamp, E. F. Stange, Differences in goblet cell differentiation between Crohn’s disease and ulcerative colitis. Differentiation. 77, 84–94 (2009).

8. D. Boltin, T. T. Perets, A. Vilkin, Y. Niv, Mucin function in inflammatory bowel disease: an update. J Clin Gastroenterol. 47, 106–111 (2013).

9. S.-J. J. Chen, X.-W. W. Liu, J.-P. P. Liu, X.-Y. Y. Yang, F.-G. G. Lu, Ulcerative colitis as a polymicrobial infection characterized by sustained broken mucus barrier. World J Gastroenterol. 20, 9468–9475 (2014).

10. B. K. Gupta, D. M. Maher, M. C. Ebeling, V. Sundram, M. D. Koch, D. W. Lynch, T. Bohlmeyer, A. Watanabe, H. Aburatani, S. E. Puumala, M. Jaggi, S. C. Chauhan, Increased expression and aberrant localization of mucin 13 in metastatic colon cancer. J Histochem Cytochem. 60, 822–831 (2012).

11. Y. H. Sheng, R. Lourie, S. K. Linden, P. L. Jeffery, D. Roche, T. V. Tran, C. W. Png, N. Waterhouse, P. Sutton, T. H. J. Florin, M. A. McGuckin, The MUC13 cell-surface mucin protects against intestinal inflammation by inhibiting epithelial cell apoptosis. Gut. 60, 1661–1670 (2011).

12. C. Moehle, N. Ackermann, T. Langmann, C. Aslanidis, A. Kel, O. Kel-Margoulis, A. Schmitz-Madry, A. Zahn, W. Stremmel, G. Schmitz, Aberrant intestinal expression and allelic variants of mucin genes associated with inflammatory bowel disease. J Mol Med (Berl). 84, 1055–1066 (2006).

13. A. Franke, D. P. B McGovern, J. C. Barrett, K. Wang, G. L. Radford-Smith, T. Ahmad, C. W. Lees, T. Balschun, J. Lee, R. Roberts, C. A. Anderson, J. C. Bis, S. Bumpstead, D. Ellinghaus, E. M. Festen, M. Georges, T. Green, T. Haritunians, L. Jostins, A. Latiano, C. G. Mathew, G. W. Montgomery, N. J. Prescott, S. Raychaudhuri, J. I. Rotter, P. Schumm, Y. Sharma, L. A. Simms, K. D. Taylor, D. Whiteman, C. Wijmenga, R. N. Baldassano, M. Barclay, T. M. Bayless, S. Brand, C. Büning, A. Cohen, J.-F. Colombel, M. Cottone, L. Stronati, T. Denson, M. De Vos, M. Dubinsky, C. Edwards, T. Florin, D. Franchimont, R. Gearry, J. Glas, A. Van Gossum, S. L. Guthery, J. Halfvarson, H. W. Verspaget, J.-P. Hugot, A. Karban, D. Laukens, I. Lawrance, M. Lemann, A. Levine, C. Libioulle, E. Louis, C. Mowat, W. Newman, J. Panés, A. Phillips, D. D. Proctor, M. Regueiro, R. Russell, P. Rutgeerts, J. Sanderson, M. Sans, F. Seibold, A. Hillary Steinhart, P. C. F Stokkers, L. Torkvist, G. Kullak-Ublick, D. Wilson, T. Walters, S. R. Targan, S. R. Brant, J. D. Rioux, R. K. Weersma, S. Kugathasan, A. M. Griffiths, J. C. Mansfield, S. Vermeire, R. H. Duerr, M. S. Silverberg, J. Satsangi, S. Schreiber, J. H. Cho, V. Annese, H. Hakonarson, M. J. Daly, M. Parkes, Genome-wide meta-analysis increases to 71 the number of confirmed Crohn’s disease susceptibility loci. Nature Publishing Group. 42 (2010), doi:10.1038/ng.717.

14. Y. H. Sheng, S. Triyana, R. Wang, I. Das, K. Gerloff, T. H. Florin, P. Sutton, M. A. McGuckin, MUC1 and MUC13 differentially regulate epithelial inflammation in response to inflammatory and infectious stimuli. Mucosal Immunol. 6, 557–568 (2013).

15. Y. H. Sheng, H. Hasnain, R. Wang, D. T. Clarke, R. Lourie, I. Oancea, K. Y. Wong, J. W. Lumley, T. H. Florin, P. Sutton, J. D. Hooper, N. A. McMillan, MUC13 protects colorectal cancer cells from death by activating the NF-κB pathway and is a potential therapeutic target - PubMed. Oncogene. 36, 700–713 (2017).

16. B. K. Gupta, D. M. Maher, M. C. Ebeling, P. D. Stephenson, S. E. Puumala, M. R. Koch, H. Aburatani, M. Jaggi, S. C. Chauhan, Functions and regulation of MUC13 mucin in colon cancer cells. J Gastroenterol. 49, 1378–1391 (2014).

17. S. Khan, M. Sikander, M. C. Ebeling, A. Ganju, S. Kumari, M. M. Yallapu, B. B. Hafeez, T. Ise, S. Nagata, N. Zafar, S. W. Behrman, J. Y. Wan, H. M. Ghimire, P. Sahay, P. Pradhan, S. C. Chauhan, M. Jaggi, MUC13 Interaction with Receptor Tyrosine Kinase HER2 Drives Pancreatic Ductal Adenocarcinoma Progression. Oncogene. 36, 491–500 (2017).

18. A. Buckley, J. R. Turner, Cell Biology of Tight Junction Barrier Regulation and Mucosal Disease. Cold Spring Harb Perspect Biol. 10, a029314 (2018).

19. A. Hartsock, W. J. Nelson, Adherens and tight junctions: structure, function and connections to the actin cytoskeleton. Biochim Biophys Acta. 1778, 660–669 (2008).

20. D. Günzel, A. S. L. Yu, Claudins and the modulation of tight junction permeability. Physiol Rev. 93, 525– 569 (2013).

21. S. Varadarajan, R. E. Stephenson, A. L. Miller, Multiscale dynamics of tight junction remodeling. J Cell Sci. 132, jcs229286 (2019).

22. J. M. Anderson, J. L. Glade, B. R. Stevenson, J. L. Boyer, M. S. Mooseker, Hepatic immunohistochemical localization of the tight junction protein ZO-1 in rat models of cholestasis. Am J Pathol. 134, 1055–1062 (1989).

23. E. Vasileva, S. Sluysmans, M.-L. Bochaton-Piallat, S. Citi, Cell-specific diversity in the expression and organization of cytoplasmic plaque proteins of apical junctions. Annals of the New York Academy of Sciences. 1405, 160–176 (2017).

24. X. Tian, Z. Liu, B. Niu, J. Zhang, T. K. Tan, S. R. Lee, Y. Zhao, D. C. H. Harris, G. Zheng, E-cadherin/β-catenin complex and the epithelial barrier. J Biomed Biotechnol. 2011, 567305 (2011).

25. H. Schmitz, C. Barmeyer, M. Fromm, N. Runkel, H. D. Foss, C. J. Bentzel, E. O. Riecken, J. D. Schulzke, Altered tight junction structure contributes to the impaired epithelial barrier function in ulcerative colitis. Gastroenterology. 116, 301–309 (1999).

26. A. J. Karayiannakis, K. N. Syrigos, J. Efstathiou, A. Valizadeh, M. Noda, R. J. Playford, W. Kmiot, M. Pignatelli, Expression of catenins and E-cadherin during epithelial restitution in inflammatory bowel disease. The Journal of Pathology. 185, 413–418 (1998).

27. N. Gassler, C. Rohr, A. Schneider, J. Kartenbeck, A. Bach, N. Obermüller, H. F. Otto, F. Autschbach, Inflammatory bowel disease is associated with changes of enterocytic junctions. Am J Physiol Gastrointest Liver Physiol. 281, G216–228 (2001).

28. C. R. Weber, S. C. Nalle, M. Tretiakova, D. T. Rubin, J. R. Turner, Claudin-1 and claudin-2 expression is elevated in inflammatory bowel disease and may contribute to early neoplastic transformation. Lab Invest. 88, 1110–1120 (2008).

29. R. J. Quin, M. A. McGuckin, Phosphorylation of the cytoplasmic domain of the MUC1 mucin correlates with changes in cell-cell adhesion. International Journal of Cancer. 87, 499–506 (2000).

30. K. Akita, M. Tanaka, S. Tanida, Y. Mori, M. Toda, H. Nakada, CA125/MUC16 interacts with Src family kinases, and over-expression of its C-terminal fragment in human epithelial cancer cells reduces cell-cell adhesion. Eur J Cell Biol. 92, 257–263 (2013).

31. C. Gao, G. Xiao, J. Hu, Regulation of Wnt/β-catenin signaling by posttranslational modifications. Cell Biosci. 4, 13 (2014).

32. X. Zhi, J. Tao, K. Xie, Y. Zhu, Z. Li, J. Tang, W. Wang, H. Xu, J. Zhang, Z. Xu, MUC4-induced nuclear translocation of β-catenin: a novel mechanism for growth, metastasis and angiogenesis in pancreatic cancer. Cancer Lett. 346, 104–113 (2014).

33. X. Huang, X. Wang, S.-M. Lu, C. Chen, J. Wang, Y.-Y. Zheng, B.-H. Ren, L. Xu, Clinicopathological and prognostic significance of MUC4 expression in cancers: evidence from meta-analysis. Int J Clin Exp Med. 8, 10274–10283 (2015).

34. L. Huang, D. Chen, D. Liu, L. Yin, S. Kharbanda, D. Kufe, MUC1 oncoprotein blocks glycogen synthase kinase 3beta-mediated phosphorylation and degradation of beta-catenin. Cancer Res. 65, 10413–10422 (2005).

35. P. Giannakouros, M. Comamala, I. Matte, C. Rancourt, A. Piché, MUC16 mucin (CA125) regulates the formation of multicellular aggregates by altering β-catenin signaling. Am J Cancer Res. 5, 219–230 (2015).

36. Y. H. Sheng, K. Y. Wong, I. Seim, R. Wang, Y. He, A. Wu, M. Patrick, R. Lourie, V. Schreiber, R. Giri, C. P. Ng, Popat, J. Hooper, G. Kijanka, T. H. Florin, J. Begun, K. J. Radford, S. Hasnain, M. A. McGuckin, MUC13 promotes the development of colitis-associated colorectal tumors via β-catenin activity. Oncogene. 38, 7294–7310 (2019).

37. L.-B. Zhou, Y.-M. Zheng, W.-J. Liao, L.-J. Song, X. Meng, X. Gong, G. Chen, W.-X. Liu, Y.-Q. Wang, D.-M. Han, N.-S. Zhong, W.-J. Lu, P.-C. Yang, X.-W. Zhang, MUC1 deficiency promotes nasal epithelial barrier dysfunction in subjects with allergic rhinitis. Journal of Allergy and Clinical Immunology. 144, 1716-1719.e5 (2019).

38. I. Gipson, S. Spurr-Michaud, A. Tisdale, Knockdown of MUC16 Alters Tight Junctions of Corneal Epithelial Cells Resulting in Decreased Transepithelial Resistance. Investigative Ophthalmology & Visual Science. 54, 554 (2013).

39. I. K. Gipson, S. Spurr-Michaud, A. Tisdale, B. B. Menon, Comparison of the transmembrane mucins MUC1 and MUC16 in epithelial barrier function. PLoS One. 9, e100393 (2014).

40. S. Resta-Lenert, S. Das, S. K. Batra, S. B. Ho, Muc17 protects intestinal epithelial cells from enteroinvasive E. coli infection by promoting epithelial barrier integrity. Am J Physiol Gastrointest Liver Physiol. 300, G1144–1155 (2011).

41. S. Meyer, M. Evers, J. H. M. Jansen, J. Buijs, B. Broek, S. E. Reitsma, P. Moerer, M. Amini, A. Kretschmer, T. ten Broeke, M. T. den Hartog, M. Rijke, C. Klein, T. Valerius, P. Boross, J. H. W. Leusen, New insights in Type I and II CD20 antibody mechanisms-of-action with a panel of novel CD20 antibodies. British Journal of Haematology. 180, 808–820 (2018).

42. S. Parry, H. S. Silverman, K. McDermott, A. Willis, M. A. Hollingsworth, A. Harris, Identification of MUC1 proteolytic cleavage sites in vivo. Biochem Biophys Res Commun. 283, 715–720 (2001).

43. J. Julian, N. Dharmaraj, D. D. Carson, MUC1 is a substrate for γ-secretase. Journal of Cellular Biochemistry. 108, 802–815 (2009).

44. R. C. Anderson, A. L. Cookson, W. C. McNabb, Z. Park, M. J. McCann, W. J. Kelly, N. C. Roy, Lactobacillus plantarum MB452 enhances the function of the intestinal barrier by increasing the expression levels of genes involved in tight junction formation. BMC Microbiol. 10, 316 (2010).

45. J. Karczewski, F. J. Troost, I. Konings, J. Dekker, M. Kleerebezem, R.-J. M. Brummer, J. M. Wells, Regulation of human epithelial tight junction proteins by Lactobacillus plantarum in vivo and protective effects on the epithelial barrier. Am J Physiol Gastrointest Liver Physiol. 298, G851–859 (2010).

46. M. Utech, R. Mennigen, M. Bruewer, Endocytosis and recycling of tight junction proteins in inflammation. J Biomed Biotechnol. 2010, 484987 (2010).

47. S. M. Stamatovic, A. M. Johnson, N. Sladojevic, R. F. Keep, A. V. Andjelkovic, Endocytosis of tight junction proteins and the regulation of degradation and recycling. Ann N Y Acad Sci. 1397, 54–65 (2017).

48. L. González-Mariscal, R. Tapia, D. Chamorro, Crosstalk of tight junction components with signaling pathways. Biochim Biophys Acta. 1778, 729–756 (2008).

49. R. O. Stuart, S. K. Nigam, Regulated assembly of tight junctions by protein kinase C. Proc Natl Acad Sci U S A. 92, 6072–6076 (1995).

50. J. Yoo, A. Nichols, J. Mammen, I. Calvo, J. C. Song, R. T. Worrell, K. Matlin, J. B. Matthews, Bryostatin-1 enhances barrier function in T84 epithelia through PKC-dependent regulation of tight junction proteins. Am J Physiol Cell Physiol. 285, C300–309 (2003).

51. J. Koizumi, T. Kojima, N. Ogasawara, R. Kamekura, M. Kurose, M. Go, A. Harimaya, M. Murata, M. Osanai, H. Chiba, T. Himi, N. Sawada, Protein kinase C enhances tight junction barrier function of human nasal epithelial cells in primary culture by transcriptional regulation. Mol Pharmacol. 74, 432–442 (2008).

52. L. W. Peterson, D. Artis, Intestinal epithelial cells: regulators of barrier function and immune homeostasis. Nat Rev Immunol. 14, 141–153 (2014).

53. G. Krause, L. Winkler, S. L. Mueller, R. F. Haseloff, J. Piontek, I. E. Blasig, Structure and function of claudins. Biochim Biophys Acta. 1778, 631–645 (2008).

54. T. Otani, M. Furuse, Tight Junction Structure and Function Revisited. Trends Cell Biol. 30, 805–817 (2020).

55. R. Al-Sadi, K. Khatib, S. Guo, D. Ye, M. Youssef, T. Ma, Occludin regulates macromolecule flux across the intestinal epithelial tight junction barrier. Am J Physiol Gastrointest Liver Physiol. 300, G1054–1064 (2011).

56. M. M. Buschmann, L. Shen, H. Rajapakse, D. R. Raleigh, Y. Wang, Y. Wang, A. Lingaraju, J. Zha, E. Abbott, E. M. McAuley, L. A. Breskin, L. Wu, K. Anderson, J. R. Turner, C. R. Weber, Occludin OCEL-domain interactions are required for maintenance and regulation of the tight junction barrier to macromolecular flux. Mol Biol Cell. 24, 3056–3068 (2013).

57. J.-C. E. Hu, F. Weiß, C. Bojarski, F. Branchi, J.-D. Schulzke, M. Fromm, S. M. Krug, Expression of tricellular tight junction proteins and the paracellular macromolecule barrier are recovered in remission of ulcerative colitis. BMC Gastroenterol. 21, 141 (2021).

58. M. Wada, A. Tamura, N. Takahashi, S. Tsukita, Loss of claudins 2 and 15 from mice causes defects in paracellular Na+ flow and nutrient transport in gut and leads to death from malnutrition. Gastroenterology. 144, 369–380 (2013).

59. P.-Y. Tsai, B. Zhang, W.-Q. He, J.-M. Zha, M. A. Odenwald, G. Singh, A. Tamura, L. Shen, A. Sailer, S. Yeruva, W.-T. Kuo, Y.-X. Fu, S. Tsukita, J. R. Turner, IL-22 Upregulates Epithelial Claudin-2 to Drive Diarrhea and Enteric Pathogen Clearance. Cell Host Microbe. 21, 671-681.e4 (2017).

60. S. Prasad, R. Mingrino, K. Kaukinen, K. L. Hayes, R. M. Powell, T. T. MacDonald, J. E. Collins, Inflammatory processes have differential effects on claudins 2, 3 and 4 in colonic epithelial cells. Lab Invest. 85, 1139– 1162 (2005).

61. F. Heller, P. Florian, C. Bojarski, J. Richter, M. Christ, B. Hillenbrand, J. Mankertz, A. H. Gitter, N. Bürgel, M. Fromm, M. Zeitz, I. Fuss, W. Strober, J. D. Schulzke, Interleukin-13 is the key effector Th2 cytokine in ulcerative colitis that affects epithelial tight junctions, apoptosis, and cell restitution. Gastroenterology. 129, 550–564 (2005).

62. S. Zeissig, N. Bürgel, D. Günzel, J. Richter, J. Mankertz, U. Wahnschaffe, A. J. Kroesen, M. Zeitz, M. Fromm, J. Schulzke, Changes in expression and distribution of claudin 2, 5 and 8 lead to discontinuous tight junctions and barrier dysfunction in active Crohn’s disease. Gut. 56, 61–72 (2007).

63. T. Oshima, H. Miwa, T. Joh, Changes in the expression of claudins in active ulcerative colitis. Journal of Gastroenterology and Hepatology. 23, S146–S150 (2008).

64. Y. Jin, A. T. Blikslager, The Regulation of Intestinal Mucosal Barrier by Myosin Light Chain Kinase/Rho Kinases. Int J Mol Sci. 21, E3550 (2020).

65. A. Banan, L. J. Zhang, M. Shaikh, J. Z. Fields, S. Choudhary, C. B. Forsyth, A. Farhadi, A. Keshavarzian, theta Isoform of protein kinase C alters barrier function in intestinal epithelium through modulation of distinct claudin isotypes: a novel mechanism for regulation of permeability. J Pharmacol Exp Ther. 313, 962–982 (2005).

66. L. González-Mariscal, E. Garay, M. Quiros, Regulation of Claudins by Posttranslational Modifications and Cell-Signaling Cascades. Current Topics in Membranes - CURR TOP MEMBR. 65, 113–150 (2010).

67. S. Aono, Y. Hirai, Phosphorylation of claudin-4 is required for tight junction formation in a human keratinocyte cell line. Exp Cell Res. 314, 3326–3339 (2008).

68. A. Y. Andreeva, E. Krause, E. C. Müller, I. E. Blasig, D. I. Utepbergenov, Protein kinase C regulates the phosphorylation and cellular localization of occludin. J Biol Chem. 276, 38480–38486 (2001).

69. T. Suzuki, B. C. Elias, A. Seth, L. Shen, J. R. Turner, F. Giorgianni, D. Desiderio, R. Guntaka, R. Rao, PKC eta regulates occludin phosphorylation and epithelial tight junction integrity. Proc Natl Acad Sci U S A. 106, 61–66 (2009).

70. E. Cario, G. Gerken, D. K. Podolsky, Toll-like receptor 2 enhances ZO-1-associated intestinal epithelial barrier integrity via protein kinase C. Gastroenterology. 127, 224–238 (2004).

71. H. Yamaguchi, T. Kojima, T. Ito, Y. Kimura, M. Imamura, S. Son, J. Koizumi, M. Murata, M. Nagayama, T. Nobuoka, S. Tanaka, K. Hirata, N. Sawada, Transcriptional Control of Tight Junction Proteins via a Protein Kinase C Signal Pathway in Human Telomerase Reverse Transcriptase-Transfected Human Pancreatic Duct Epithelial Cells. The American Journal of Pathology. 177, 698–712 (2010).

72. D. Iitaka, S. Moodley, H. Shimizu, X.-H. Bai, M. Liu, PKCδ–iPLA2–PGE2–PPARγ signaling cascade mediates TNF-α induced Claudin 1 expression in human lung carcinoma cells. Cellular Signalling. 27, 568–577 (2015).

73. T. Breugelmans, H. Van Spaendonk, J. G. De Man, H. U. De Schepper, A. Jauregui-Amezaga, E. Macken, S. K. Lindén, I. Pintelon, J.-P. Timmermans, B. Y. De Winter, A. Smet, In-Depth Study of Transmembrane Mucins in Association with Intestinal Barrier Dysfunction During the Course of T Cell Transfer and DSS-Induced Colitis. J Crohns Colitis. 14, 974–994 (2020).

74. S. J. Williams, D. H. Wreschner, M. Tran, H. J. Eyre, G. R. Sutherland, M. A. McGuckin, Muc13, a novel human cell surface mucin expressed by epithelial and hemopoietic cells. J Biol Chem. 276, 18327–18336 (2001).

75. Y. H. Sheng, S. Triyana, R. Wang, I. Das, K. Gerloff, T. H. Florin, P. Sutton, M. A. McGuckin, MUC1 and MUC13 differentially regulate epithelial inflammation in response to inflammatory and infectious stimuli. Mucosal Immunol. 6, 557–568 (2013).

76. R. Elmentaite, N. Kumasaka, K. Roberts, A. Fleming, E. Dann, H. W. King, V. Kleshchevnikov, M. Dabrowska, S. Pritchard, L. Bolt, S. F. Vieira, L. Mamanova, N. Huang, F. Perrone, I. Goh Kai’En, S. N. Lisgo, M. Katan, S. Leonard, T. R. W. Oliver, C. E. Hook, K. Nayak, L. S. Campos, C. Domínguez Conde, E. Stephenson, J. Engelbert, R. A. Botting, K. Polanski, S. van Dongen, M. Patel, M. D. Morgan, J. C. Marioni, O. A. Bayraktar, K. B. Meyer, X. He, R. A. Barker, H. H. Uhlig, K. T. Mahbubani, K. Saeb-Parsy, M. Zilbauer, M. R. Clatworthy, M. Haniffa, K. R. James, S. A. Teichmann, Cells of the human intestinal tract mapped across space and time. Nature. 597, 250–255 (2021).

77. V. Solovyev, “V. Solovyev, A Salamov (2011) Automatic Annotation of Microbial Genomes and Metagenomic Sequences. In Metagenomics and its Applications in Agriculture, Biomedicine and Environmental Studies (Ed. R.W. Li), Nova Science Publishers, p.61-78.” in (2011), pp. 61–78.

78. X. Li, R. W. Wubbolts, N. M. C. Bleumink-Pluym, J. P. M. van Putten, K. Strijbis, The Transmembrane Mucin MUC1 Facilitates β1-Integrin-Mediated Bacterial Invasion. mBio. 12, e03491–20 (2021).

